# Heterogeneity in the *M. tuberculosis* β-lactamase inhibition by Sulbactam

**DOI:** 10.1101/2022.12.06.519319

**Authors:** Tek Narsingh Malla, Kara Zielinski, Luis Aldama, Sasa Bajt, Denisse Feliz, Brendon Hayes, Mark Hunter, Christopher Kupitz, Stella Lisova, Juraj Knoska, Jose Martin-Garcia, Valerio Mariani, Suraj Pandey, Ishwor Poudyal, Raymond G. Sierra, Alexandra Tolstikova, Oleksandr Yefanov, Chung Hong Yoon, Abbas Ourmazd, Petra Fromme, Peter Schwander, Anton Barty, Henry N. Chapman, Emina A. Stojkovic, Alex Batyuk, Sébastien Boutet, George N. Phillips, Lois Pollack, Marius Schmidt

**Affiliations:** Physics Department, University of Wisconsin-Milwaukee, 3135 N Maryland Ave, Milwaukee, WI 53211, USA; School of Applied and Engineering Physics, Cornell University, 254 Clark Hall, Ithaca, NY 14853, USA; Department of Biology, Northeastern Illinois University, 5500 N. St. Louis Ave., Chicago, Illinois 60625, USA; The Hamburg Centre for Ultrafast Imaging, Luruper Chaussee 149, 22761 Hamburg, Germany; Center for Free-Electron Laser Science CFEL, Deutsches Elektronen Synchrotron, Notkestrasse 85, 18 22607 Hamburg, Germany; Linac Coherent Light Source LCLS, SLAC National Accelerator Laboratory, 2575 Sand Hill Road, Menlo Park, CA 94025, USA; Department of Crystallography and Structural Biology, Institute of Physical Chemistry, Rocasolano, Spanish National Research Council (CSIC), Serrano 119, 28006 Madrid, Spain; Deutsches Elektronen-Synchrotron DESY, Notkestrasse 85, 22607 Hamburg, Germany; School of Molecular Sciences and Biodesign Center for Applied Structural Discovery, 20 Arizona State University, Tempe, AZ 85287-1604, USA; Center for Data and Computing in Natural Science CDCS, Deutsches Elektronen-Synchrotron DESY, Notkestrasse 85, 22607 Hamburg, Germany; Department of Physics, Universität Hamburg, Luruper Chaussee 149, 22761 Hamburg, Germany; Department of BioSciences, Rice University, 6100 Main Street, Houston, Texas 77005, USA; Department of Chemistry, Rice University, 6100 Main Street, Houston, Texas 77005, USA

## Abstract

For decades, researchers have been determined to elucidate essential enzymatic functions on the atomic lengths scale by tracing atomic positions in real time. Our work builds on new possibilities unleashed by mix-and-inject serial crystallography (MISC)^1-5^ at X-ray free electron laser facilities. In this approach, enzymatic reactions are triggered by mixing substrate or ligand solutions with enzyme microcrystals^6^. Here, we report in atomic detail and with millisecond time-resolution how the *Mycobacterium tuberculosis* enzyme BlaC is inhibited by sulbactam (SUB). Our results reveal ligand binding heterogeneity, ligand gating^7-9^, cooperativity, induced fit^10,11^ and conformational selection^11-13^ all from the same set of MISC data, detailing how SUB approaches the catalytic clefts and binds to the enzyme non-covalently before reacting to a *trans-*enamine. This was made possible in part by the application of the singular value decomposition^14^ to the MISC data using a newly developed program that remains functional even if unit cell parameters change during the reaction.

## 1. Introduction

Beta(β)-lactamases are bacterial enzymes that provide multi-resistance to β-lactam antibiotics. They inactivate the β-lactam antibiotics by hydrolyzing the amide bond of the β-lactam ring^15,16^ (Fig. 1 a). Our study focuses on β-lactamase from *Mycobacterium tuberculosis* (*Mtb*)-the causative agent of tuberculosis. *Mtb* β-lactamase (BlaC) is a broad-spectrum Ambler class A^17^ β-lactamase capable of hydrolyzing all classes of β-lactam antibiotics used for treatment of tuberculosis ^18,19^. In the latest report, the World Health Organization warned that multidrug-resistant tuberculosis is a public health crisis and a health security threat. In 2020, Tuberculosis was the 13th leading cause of death and the second leading infectious killer after COVID-19 (above HIV/AIDS), claiming 1.5 million lives worldwide ^20^. The rapid and worldwide emergence of antibiotic resistant bacteria, including *Mtb*, is endangering the efficacy of antibiotics, causing the Center for Disease Control (CDC) to classify several bacteria as urgent and serious threats ^21^. Research efforts focused on deciphering the molecular mechanism of antibiotic resistance within microbial pathogens such as *Mtb* can aid in novel-drug design, resistant to β-lactamase action, and therefore, contribute to managing this evolving crisis.

**Figure 1.**
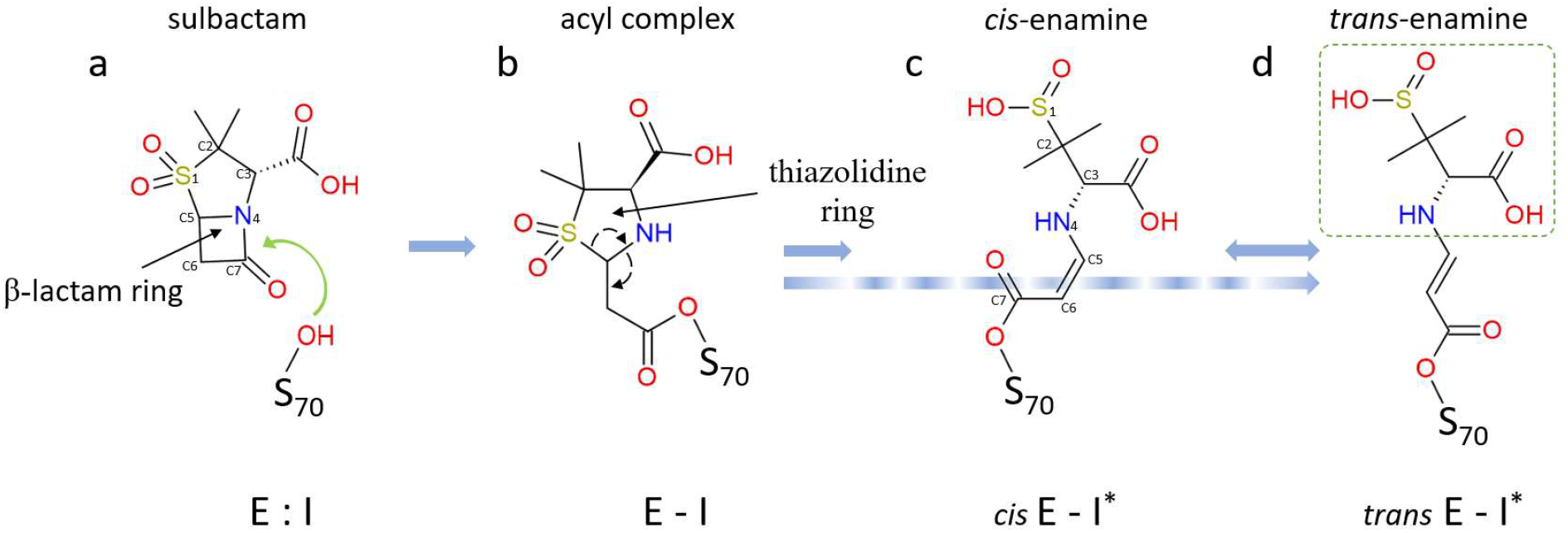
Simplified mechanism of sulbactam inhibition reaction of BlaC. (a) Non-covalent enzyme-inhibitor complex formation (E:I). The characteristic β-lactam ring is marked. The nucleophilic attack by Ser70 is shown by a green arrow. (b) The nucleophilic attack opens the β-lactam ring, and a covalent bond is created between Ser70 and sulbactam resulting in the acyl-enzyme complex (E-I). In this current state, the E-I is very unstable and causes reorganization of bonds shown by dotted arrows. This modification leads to permanent opening of the 5-member thiazolidine ring and formation of either (c) *cis-*enamine (*cis* E-I^*^) and then to (d) *trans-*enamine (*trans* E-I^*^) or directly to *trans-*enamine following the blue dotted arrow. *Cis-* and *trans-*enamines differ in the configuration of the C5=C6 double bond. A second nucleophilic attack by Ser128 on C5 may lead to cleavage of the fragment shown by dotted box in (d). Further modifications are possible which are not shown here.

Structural information involving enzyme-substrate reaction intermediates is essential for understanding the mechanism of enzymes in action. Although several static X-ray structures of BlaC with various substrates have been determined ^18,22-27^, the reaction intermediates of β-lactamase reaction involving different antibiotic substrates remain mostly unknown. With time-resolved crystallography (TRX), conformational changes in biological macromolecules can be explored in real time ^28^. Once a reaction is initiated within the crystals, ensemble-averaged structures are obtained along a reaction pathway. In addition, the chemical kinetics that governs the biological reaction can be deduced ^14,29-32^. With mix-and-inject serial crystallography (MISC)^1^ single-turnover enzymatic reactions can be investigated in a time resolved manner. The substrate solution is mixed with enzyme micro-crystals before the mixture is injected into the X-ray beam. The reaction is triggered by diffusion of substrate and the resulting change is probed after a delay by short X-ray pulses ^3,4,33^. Using MISC, intermediates within the BlaC reaction with the third-generation cephalosporin-based antibiotic, ceftriaxone (CEF) were previously characterized from 5 ms to 2s ^1,33,34^.

Irreversible enzyme inhibitors represent potential new drugs in the fight against antibiotic resistance. Sulbactam, clavulanate and tazobactam, all suicide inhibitors with a β-lactam ring, irreversibly bind to β-lactamases and block their activity ^35,36^. As a result, the β-lactam antibiotics are protected from enzymatic degradation, which helps retain their efficacy. Due to its excellent solubility, sulbactam (SUB) is a superb candidate for a MISC experiments involving BlaC.

Raman microscopy and mass spectrometry suggest a simplified mechanism for SUB binding to BlaC ^37,38^, shown in Fig. 1. The first step is the formation of a reversible (non-covalent) enzyme inhibitor complex in the active site (E:I). Following this association step, the nucleophilic attack by the catalytically active serine on the β-lactam ring of the inhibitor leads to the formation of a short-lived, covalently bound acyl-enzyme intermediate (E-I). Further modifications of the chemical structure of the inhibitor lead to an inactivated enzyme (E-I^*^). The structures of intermediates shown in Fig. 1 are relevant to the observed results presented in this paper within the measured timescale. SUB has also been described as a substrate of class A β-lactamases and can be hydrolyzed albeit at much slower rate than β-lactam antibiotics ^35,39^.

BlaC can be crystallized in a monoclinic space group with four subunits A - D in the asymmetric unit (Fig. 2 a) ^1,40^. In this crystal form, the BlaC structure displays a large cavity of 30 Å diameter in the center. Additional cavities as large as 90 Å are identified within adjacent asymmetric units that allow easy diffusion of ligand molecules through the crystalline lattice. The E-I^*^ structure of the BlaC-SUB has been determined at cryogenic temperature after soaking BlaC crystals for several minutes with SUB ^27,41-43^. More recently, Pandey and colleagues captured an intermediate (presumably E:I) at a single time point (66 ms) after mixing BlaC crystals with SUB ^34^. Although subunits B/D displayed already a covalently bound adduct, an intact, non-covalently bound SUB was observed in subunits A/C (Fig. 2 a). Subunits A/C are rather inactive, since they did not participate at all in the reaction with CEF in earlier experiments ^1,33^. Given the inactivity of subunits A/C, it was not clear whether the reaction with SUB takes more time to complete or proceed at all. Therefore, a time-series of MISC datasets is necessary, to further investigate both the binding of SUB inhibitor to and its subsequent reaction with the BlaC.

**Figure 2.**
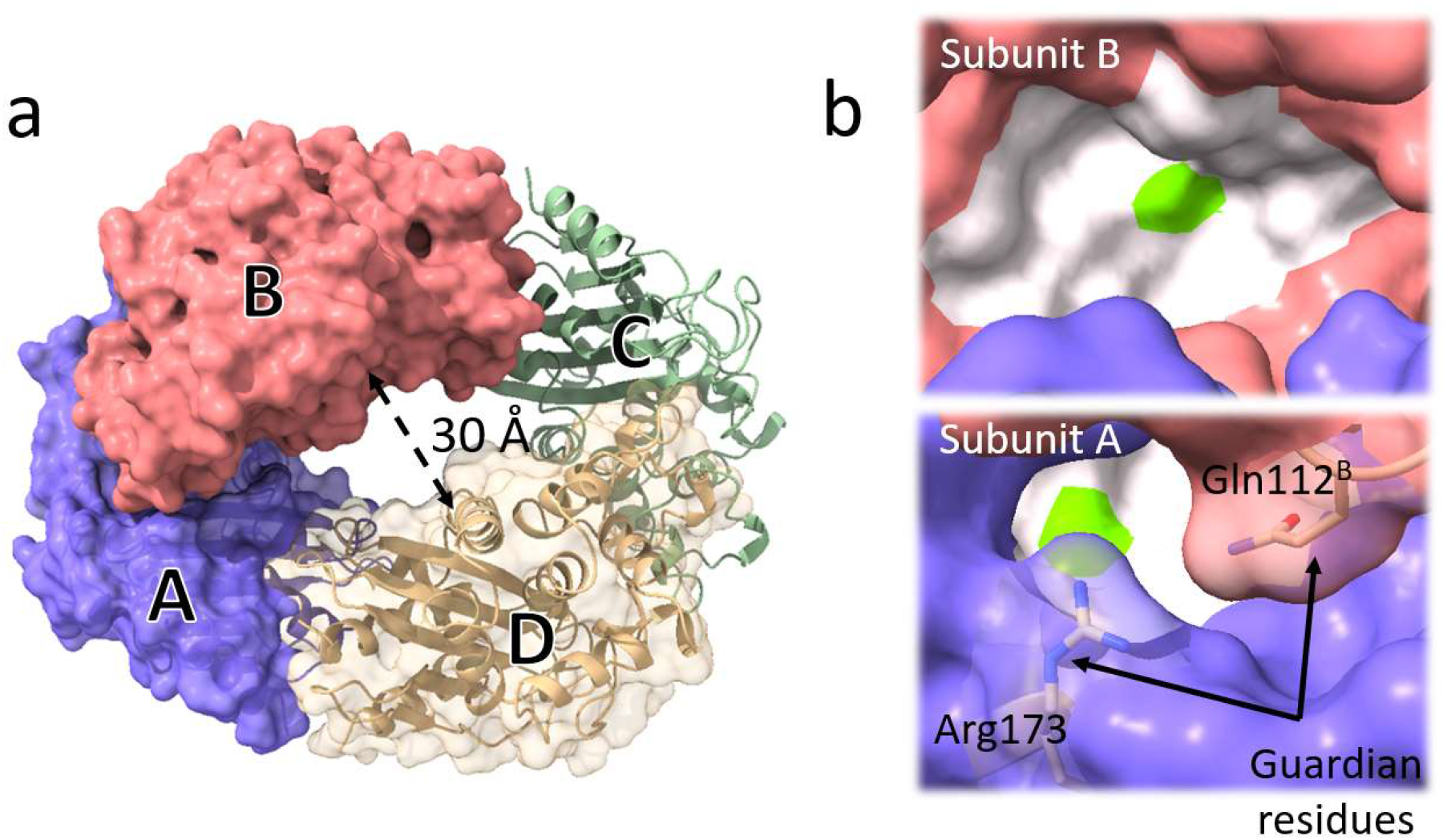
Structure of BlaC. (a) Subunits A – D in the asymmetric unit are marked and shown by blue, red, green, and yellow respectively. (b) Binding pockets of subunit B (top) and subunit A (bottom). The active site is represented by the white surface. The position of the catalytically active serine is marked in green. The access to the active site in subunit B is wide open. The entrance to the active site of subunit A is partially occluded by two residues (Glutamine and Arginine) called the guardian residues.

As in any time-resolved experiment, except in those performed on ultrafast time scales ^44-47^, multiple states can mix into any single time point observed during the reaction ^48^. As demonstrated for X-ray data ^14^ these mixtures can be characterized and potentially separated using the singular value decomposition (SVD). Within this context, SVD is an unsupervised machine learning algorithm ^49^ that can inform from time-resolved X-ray data the number of observable processes which is equivalent to the number of relaxation times and the number of structurally distinguishable time-independent reaction intermediates ^14^. In addition, it can provide information regarding the kinetic mechanism and the energetics of the reaction ^50-54^. SVD has never been applied to a time-series from a MISC experiment. Originally, the SVD method was developed to work with isomorphous difference maps assuming that the unit cells in the crystals do not change during a reaction. However, the unit cell parameters of the BlaC crystals vary after mixing (Tab. 1). This makes a data analysis approach that relies on a stable volume of interest very challenging. Therefore, a new suite of programs “pySVD4TX” was developed, that remains functional even when the unit cell parameters change.

**Table 1.** Data collection and refinement statistics

|  | Water | 3ms | 6ms | 15ms | 30ms | 240ms | 700ms | 3 h soak |
| --- | --- | --- | --- | --- | --- | --- | --- | --- |
| Space Group | P2<sub>1</sub> |  |  |  |  |  |  |  |
| a,b,c (Å),<br>β (°) | 79.6,98.1,<br>110.9,108.5 | 80.5,98.8,<br>111.5,108.5 | 80.4,8.8,<br>112.1,108.7 | 80.3,98.6,<br>112.1,108.8 | 80.2,98.5,<br>112.3,108.9 | 79.8,97.9,<br>113.2,109.3 | 80.0,98.3,<br>112.1,108.8 | 79.4, 96.7,<br>111.1,108.5 |
| Resolution<br>range (Å) | 22.14-2.2<br>(2.3 -2.2)<sup>†</sup> | 22.01-2.6<br>(2.75-2.6) | 20.53-2.75<br>(2.85-2.75) | 20.86-2.65<br>(2.75-2.65) | 20.4-2.95<br>(3.06-2.95) | 22.42-2.35<br>(2.43-2.34) | 20.94-3.2<br>(3.38-3.2) | 22.3-2.7<br>(2.8-2.7) |
| Hits | 88,978 | 61,508 | 26,309 | 51,166 | 27,143 | 75,038 | 2,854 |  |
| Indexed<br>patterns | 73,965 | 36,400 | 17,596 | 31,958 | 11,123 | 59,928 | 2,740 |  |
| Observed<br>Reflections | 37,857,581 | 14,718,802 | 6,258,604 | 11,092,596 | 3,888,789 | 29,990,504 | 1,582,566 |  |
| Unique<br>reflections | 83,447 | 42,822 | 43,284 | 40,586 | 32,517 | 68,441 | 27,514 | 38,773 |
| Multiplicity | 453(335) | 343(164) | 144(69) | 273(130) | 119(67) | 446(306) | 57(35) | 79(28) |
| Completeness | 100(100) | 100(100) | 100(100) | 100(100) | 100(100) | 100(100) | 99.9(99.9) | 100(100) |
| R<sub>split/merge</sub> (%) | 23.3(528) | 23.4(427) | 24.3(354) | 23.6(316) | 30.4(176) | 28.8(689) | 44.6(208) | 18.2(803) |
| CC* (%) | 99.6(69.8) | 99.3(55.9) | 99.0(51.6) | 98.9(58.7) | 98.2(52.9) | 99.3(64.2) | 95.9(51.3) | 99.6(64.2) |
| Resolution<br>range | 22.14-2.2 | 22.01-2.6 | 20.53-2.75 | 20.86-2.65 | 20.4-2.95 | 22.42-2.35 | 20.94-3.2 | 22.3-2.6 |
| Reflections<br>used | 68064 | 40968 | 41929 | 38742 | 33082 | 53680 | 27127 | 25,400 |
| R<sub>cryst</sub>/R<sub>free</sub> | 0.22/0.25 | 0.22/0.24 | 0.22/0.24 | 0.23/0.26 | 0.22/0.26 | 0.23/0.25 | 0.27/0.29 | 0.22/0.27 |
| Occupancy<br>(in %) | n.a | n.a. | n.a. | TEN<sup>B</sup>: 96<br>TEN<sup>D</sup>: 90 | SUB<sup>A</sup>: 52<br>TEN<sup>B</sup>: 85<br>SUB<sup>C</sup>: 63<br>TEN<sup>D</sup>: 77 | TEN<sup>A</sup>: 100<br>TEN<sup>B</sup>: 100<br>TEN<sup>C</sup>: 100<br>TEN<sup>D</sup>: 100 | TEN<sup>A</sup>:100<br>TEN<sup>B</sup>:100<br>TEN<sup>C</sup>:100<br>TEN<sup>D</sup>:100 | TEN<sup>A</sup>:98<br>TEN<sup>B</sup>:96<br>TEN<sup>C</sup>:98<br>TEN<sup>D</sup>:88 |
| r.m.s.d<sup>‡</sup> bond<br>length | 0.004 | 0.002 | 0.003 | 0.003 | 0.003 | 0.003 | 0.002 | 0.003 |
| r.m.s.d bond<br>angles | 0.641 | 0.545 | 0.601 | 0.642 | 0.768 | 0.65 | 0.428 | 0.584 |
| No. of H<sub>2</sub>O | 447 | 140 | 203 | 183 | 44 | 311 | 16 | 90 |
| <sup>†</sup> Values in parentheses represent the highest resolution bin<br><sup>A,B,C,D</sup> The superscript letters represent corresponding subunits. |  |  |  |  |  |  |  |  |

## Results

### 4.1 Binding of sulbactam

The four subunits of BlaC found in the asymmetric unit arrange in shape reminiscent to a torus (Fig. 2 a). The alternating subunits are display similar binding kinetics while significantly differing from the adjacent ones. Here, subunits A and C share similarities. As do subunits B and D. However, there are distinct differences between these pairs. Whereas the catalytic clefts of subunits B and D are wide open, those of subunits A and C are partially occluded by the neighboring subunits (Fig. 2 b). Particularly the residues Gln109^B/D^ and Gln108^B/D^ (the superscript B/D denotes residues from the neighboring subunits B and D, respectively) prevent substrate diffusion from the center of the torus, and Gln112^B/D^ and Arg173 block access to the active site from the exterior. The reaction of SUB with BlaC was followed by difference electron density (DED) maps obtained at MISC delays (Δt_misc_) of 3 ms, 6 ms, 15 ms, 30 ms, 270 ms and 700 ms. At 3, 6 and 15 ms the DED features near the active site of subunit A are weak (Fig. 3 a-c). Substantial displacements of the residues flanking the active site are observed (Tab. 2). Particularly, the long side chain residues like Gln112^B^ and Arg173 (called here the guardian residues, Fig. 2 b) move outward of the active site before relaxing back to original positions. Next to these residues at 3ms, almost 10 Å away from Ser70, there are positive densities that are spatially more spread out than that of water. A SUB molecule can be placed in the electron density (Fig. 3 a). However, after refinement negative density (red contour lines) appear in the F_obs_-F_calc_ map (Fig. 4). In addition, the B-factor values of the SUB atoms are all in excess of 120 Å^2^. The translational and rotational disorder of the free SUB that accumulates near the guardian residues renders the refinement of a single conformation difficult. At 6 and 15ms, density appear closer towards Ser70 (Fig. 3 b-c). These could indicate an initial trace of SUB molecules migrating to the active site after being held up by the guardian residues. Up to 15 ms, these densities are too weak that a SUB molecule can be placed with confidence.

**Figure 3.**
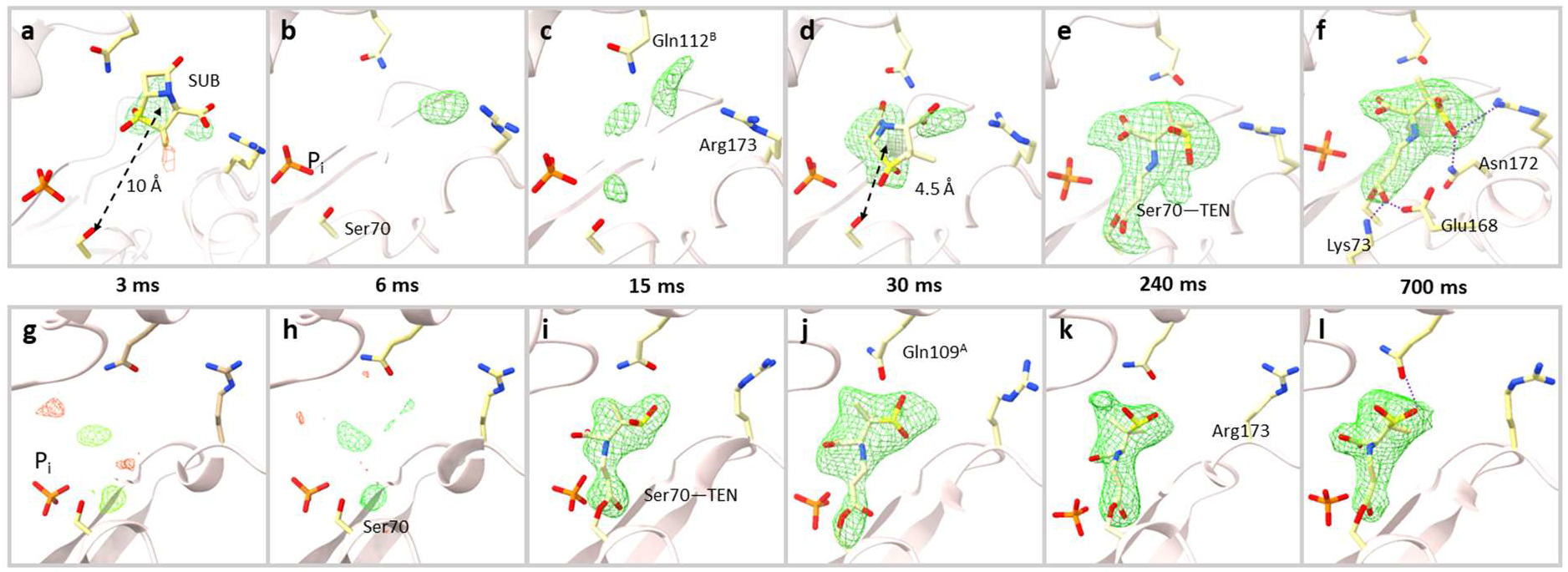
Difference electron density maps at the active site of subunit A and subunit B (contour level ±3σ). *Subunit A, top row:* (a) At 3ms, weak densities can be identified at the entrance of cavity between Gln112^B^ and Arg173 and a SUB placed there. (b) At 6ms, very weak density is observed. Residues Gln112^B^ and Arg173 continue to move. The phosphate molecule (P_i_) near the active site is marked. (c) At 15ms, difference density features are identified closer to the catalytically active residue Ser70. The guardian residues (Gln112^B^ and Arg173) that are located at the entrance to the binding pocket are marked. (d) At 30 ms a strong DED feature appears within the active site. An intact SUB molecule is placed there. (e) At 240 ms, the SUB has reacted with Ser70 to form TEN giving rise to an elongated density. (f) At 700 ms, the elongated density of the TEN is fully developed. Additional hydrogen bonds between the TEN and other side chains are shown. *Subunit B, bottom row:* (g-h) At 3 and 6ms, no interpretable density was present in the catalytic center. (i) At 15ms, the SUB has already reacted with Ser70 to from TEN. (j-l) TEN densities as observed at Δ_misc_ from 30 ms to 700 ms. Gln109^A^ and Arg173 are marked in j and k, respectively.

**Table 2.** Important distances (in Å) in the active centers of subunit A and B. Values in parentheses represent the change in the position [in Å] of the terminal side chain atom from its original position at 0 ms.

|  | 0 ms | 3ms | 6ms | 15ms | 30ms | 240ms | 700ms |
| --- | --- | --- | --- | --- | --- | --- | --- |
| (a) Distance between side chain of some active residues in subunit A |  |  |  |  |  |  |  |
| Ser70 OG to Arg173 CZ | 10.9 | 11.6<br>(2.7) | 12.1<br>(3.1) | 10.7<br>(1.2) | 11.0<br>(1.6) | 10.4<br>(1.4) | 10.2<br>(1.3) |
| Ser70 OG to Gln112<sup>B</sup> CD | 10.6 | 12.0<br>(2.2) | 10.9<br>(0.4) | 10.9<br>(1.3) | 10.6<br>(0.4) | 9.9<br>(0.5) | 9.8<br>(0.2) |
| Ser70 OG to Gln109<sup>B</sup> CD | 9.0 | 10.2 | 10.0 | 10.3 | 10.2 | 8.6 | 8.7 |
| (b) Distance between side chains of some active residues in subunit B |  |  |  |  |  |  |  |
| Ser70 OG to Arg173 CZ | 13.9 | 13.2<br>(2.1) | 13.9<br>(1.7) | 14.4<br>(1.6) | 14.8<br>(0.9) | 14.9<br>(1.5) | 14.4<br>(1.2) |
| (c) Distances from BlaC side chains to selected SUB and TEN atoms in subunit A |  |  |  |  |  |  |  |
| Thr239 O to SUB O12 | - | - | - | - | 2.7 | - | - |
| Gln112<sup>B</sup> NE2 to SUB O8 | - | - | - | - | 2.7 | - | - |
| Lys73 NZ to TEN O8 | - | - | - | - | - | 2.5 | 3.1 |
| Glu168 OE2 to TEN O8 | - | - | - | - | - | 2.6 | 2.5 |
| Asn172 ND2 to TEN O12 | - | - | - | - | - | 2.5 | 2.9 |
| (d) Distances from BlaC side chains to selected TEN atoms in subunit B |  |  |  |  |  |  |  |
| Thr239 O to TEN O8 | - | - | - | 2.9 | 3 | 3.1 | 3.2 |
| Gln109<sup>A</sup> NE2 to TEN O12 | - | - | - | 3.9 | 3.4 | 3.2 | 4.1 |
| <sup>A,B</sup> subunits that host the amino acid. |  |  |  |  |  |  |  |

**Figure 4.**
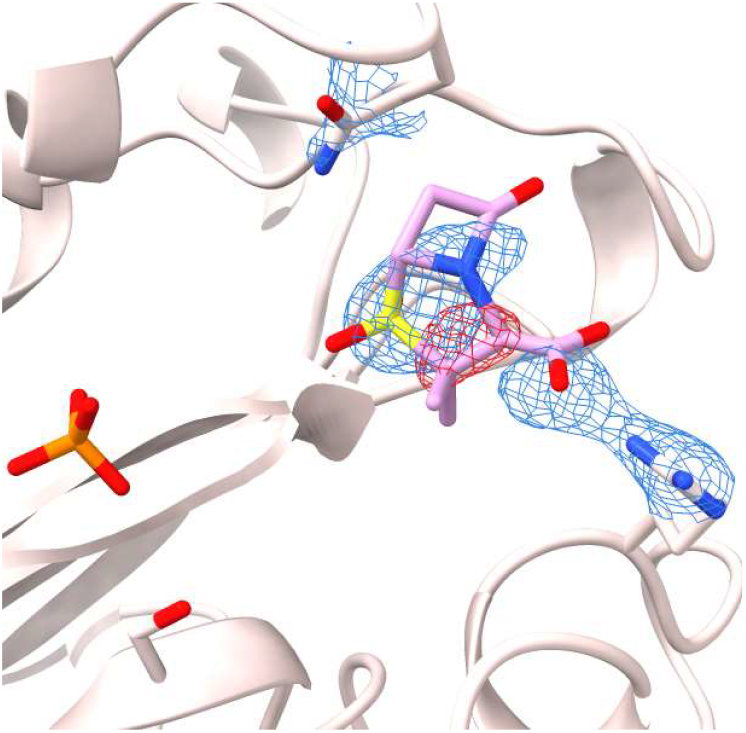
The appearance of SUB at the entrance to the active site. SUB (purple) was placed in the DED_omit_ feature near the guardian residues observed at Δt_MISC_=3 ms (Fig. 3 a) and refined.

At 30 ms, stronger DED features (max σ = 5.5) appear around 4.5 Å from the catalytically active Ser70 (Fig. 3 d). An intact sulbactam can be modelled which reproduces and corroborates the findings at 66 ms obtained from an earlier experiment ^34^. The β-lactam ring is oriented away from the Ser70. Between 66 ms and 240 ms, the SUB must have rotated so that the β-lactam ring is positioned towards the Ser70 at which point the nucleophilic attack occurs (Fig. 1 a). At 240 ms, the elongated DED feature that originates from the Ser70 directly supports the presence of a covalently bound *trans*-enamine (TEN) (Fig. 3 e). The BlaC-TEN adduct structurally relaxes until 700 ms, the final time point in the time series (Fig. 3 f). These snapshots of the reaction in progress were assembled to a movie of an enzyme in action.

In subunit B, there is no evidence of an intact SUB that accumulates in the active site (Fig. 3 g-h). Even the features that appeared near Arg173 in subunits A/C are not present. However, at 15 ms, the presence of strong DED that extends from the Ser70 supports a covalently bound TEN (Fig. 2 b, Fig. 3 i). The occupancy refined to 85% in B and 86% in D. This suggests the reaction is close to completion. At later time points, no large changes in the structure of the BlaC-TEN complex are apparent. (Fig. 3 j-l). However, in subunits A/C, the reaction keeps progressing and structural changes of the TEN are still observed (Tab. 2)

### 4.3 Reaction of BlaC after treatment with SUB for a longer period of time

To investigate SUB binding to BlaC on a longer time scale, static cryo structure was determined by soaking BlaC macrocrystals in SUB solution for 3 hours. *Trans-*enamines were identified in all subunits with close to 100% occupancy (Fig. 5).

**Figure 5.**
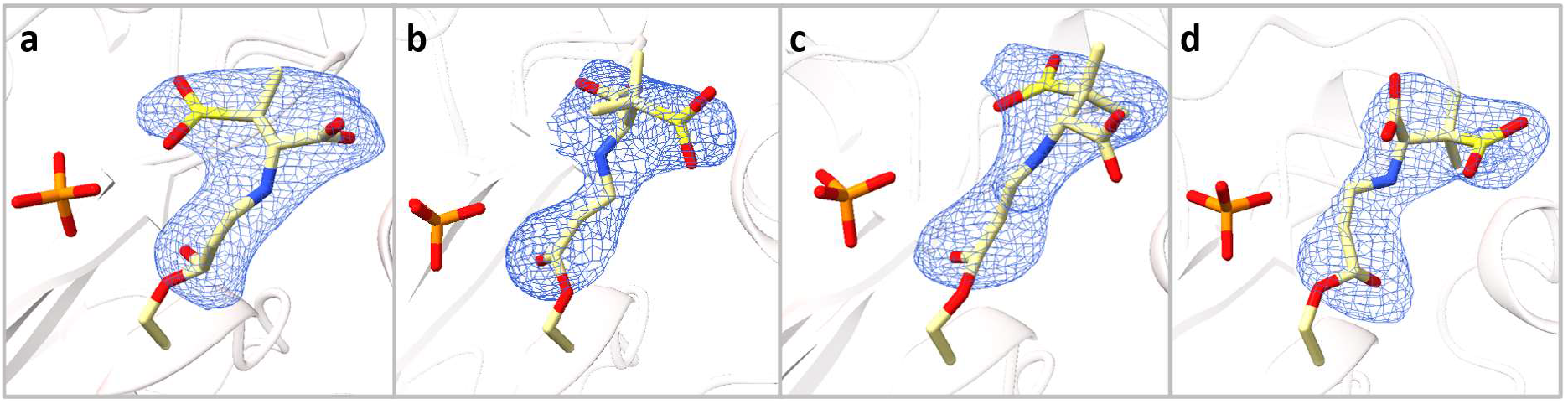
Difference electron density maps determined in the active sites of BlaC after 3 h of soaking with SUB. (a-d) Polder omit maps in each subunit of BlaC contoured at 2.5 σ. All the species are characterized by covalently bound *trans-*enamines.

### 4.3 Temporal Variation of Difference Electron Density

The SVD analysis is required to identify the number of intermediates as well as the relaxation times from the time-series of DED maps ^14^ (see Methods). The right singular vectors (rSVs) obtained from the SVD analysis plotted as a function of MISC time delays represent the temporal variation of the reaction. It is important that the relaxation processes inherent to each rSV are accurately determined. The slow initial progress and sudden increase of the magnitude of DED values in the active sites require an appropriate function that could account for this behavior. An excellent fit was obtained by Eqn. 2 which consists of a (step-like) logistic function that accounts for the steep first phase and an exponential saturation component. Detailed values of the parameters of Eqn. 2 are shown in the Tab. 3.

**Table 3.** Relaxation times and amplitudes determined from fitting Eqn. 2 to the significant rSVs.

|  | Subunit A | Subunit B | Subunit C | Subunit D |
| --- | --- | --- | --- | --- |
| $\lambda$ (ms<sup>-1</sup>) | 0.25 | 0.51 | 0.33 | 0.69 |
| $\tau_1$ (ms) | 23.26 | 10.17 | 25.53 | 9.81 |
| $\tau_2$ (ms) | 86.03 | 80.41 | 90.01 | 88.15 |
| **rSV<sub>1</sub>** |  |  |  |  |
| A<sub>0,1</sub> | 11.72 | 38.32 | 8.89 | 32.77 |
| A<sub>1,1</sub> | 118.95 | 195.44 | 113.3 | 155.37 |
| A<sub>2,1</sub> | 134.48 | 122.15 | 152.08 | 137.01 |
| **rSV<sub>2</sub>** |  |  |  |  |
| A<sub>0,2</sub> | -17.96 | -0.25 | -12.95 | 3.23 |
| A<sub>1,2</sub> | -90.67 | -41.79 | -74.92 | 26.54 |
| A<sub>2,2</sub> | 142.88 | 64.27 | 117.31 | -48.72 |
| $\lambda$, $\tau_1$ and $\tau_2$ are global fit parameters for all significant rSVs; the amplitudes A's are determined independently for each rSV. | | | | |

The first two significant rSVs for each subunit are shown by blue and red squares respectively (Fig. 6 a-d). They allow for the determination of relaxation times τ_1_ and τ_2_ by fitting Eqn. 2, shown by their respective colored solid line. The rest of the rSVs, shown by colored diamonds in Fig. 6 a, are distributed closely around zero and do not contribute to the subsequent analysis. The process with relaxation time τ_1_ results from the first appearance of DED in the active sites. In subunits A/C, this process reflects the appearance of the non-covalently bound SUB which occurs at around 23.5 and 24.5 ms, respectively. In subunits B/D, a covalently bound TEN is observed at 9.6 ms and 10.2 ms respectively after mixing. In subunits A/C, the process τ_2_ results from the transformation of intact SUB to covalently bound TEN which happens approximately 75 ms and 90 ms after mixing. In contrast, in subunits B/D TEN is present already during the first phase (Fig. 3 g-l; Fig 2 e-h) and no further chemical modification of the TEN is observed. Despite this, a second relaxation process is also observed which coincides with the SUB to TEN formation in subunits A/C.

**Figure 6.**
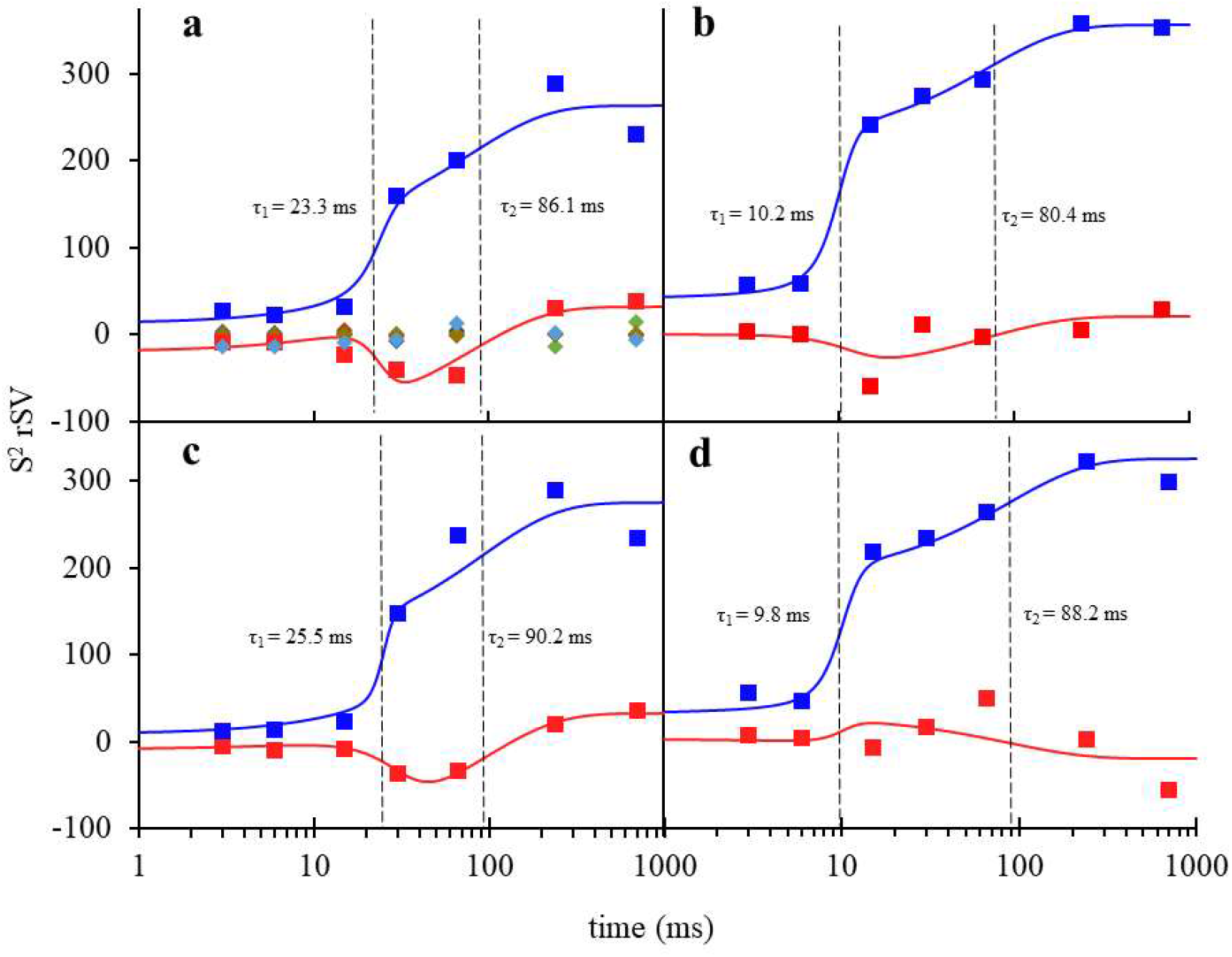
Right singular vectors (rSVs) derived from the MISC time-courses of DED maps in the active sites of the BlaC. (a) Right singular vectors plotted as a function of Δt_MISC_ for subunits A. The first and second significant right singular vectors are shown by blue and red triangles respectively. Solid colored lines are the result of a global fit of Eqn. 2 to the significant rSVs. The colored diamonds represent insignificant rSVs derived from subunit A. (b-d) Significant rSVs plotted as a function of Δt_MISC_ for subunits B, C and D respectively. Colors and lines as in panel (a). The vertical dashed black lines in all panels denote the relaxation times τ_1_ and τ_2_ that result from the fit. For subunits A and C, τ_1_ belongs to accumulation of SUB and τ_2_ corresponds to formation of covalently bound TEN from intact SUB. For subunits B and D, τ_1_ is the time when direct formation of TEN occurs and τ_2_ belongs to a second phase of relaxation.

### 4.4 Inhibitor Diffusion

The kinetics of SUB binding to subunits B/D is evaluated first, since it allows for an estimate of the ligand concentration in the unit cell that is required to analyze the observations for subunits A/C. The total concentration of BlaC monomers in the crystals (E_free_) is ∼16 mM. If only subunit B is considered, the effective [E_free_] is 4 mM. The apparent diffusion time of SUB into the microcrystals is ∼7 ms (Tab. 4). It needs to be pointed out that MISC does not directly measure diffusion inside crystals. Instead, free ligand concentrations, and related to them the diffusion time, are estimated only indirectly through the rate equations (Eqns. 3 and 4) which ultimately must reproduce the observed (refined) ligand occupancies ^34^. The unknown parameters in these equations must be varied until the concentration of species [E:I] and [E-I^*^] best match their respective occupancy values (Fig. 7 a). Additionally, relaxation times derived from the concentration profiles must also agree with those obtained from SVD analysis of the DED maps (compare Tabs. 1 and 4).

**Table 4.** Characterization of SUB diffusion and reaction rate coefficients

| (a) Parameters to estimate the diffusion of SUB into the active sites (Eqns. 3, 4 and 6). |  |  |  |  |
| --- | --- | --- | --- | --- |
| $\mu$ | 0.59 ms<sup>-1</sup> | | | |
| $t_0$ | 7 ms | | | |
| $\Delta I_0$ | 25 mM | | | |
| (b) Rate coefficients and approximate characteristic times (estimated from the half maximum of the blue, orange and green curves shown in Figs. 10 and 12). |  |  |  |  |
|  | Subunits A and C |  | Subunits B and D |  |
| Rate Coefficients<sup>a</sup> | $k_{\text{max,entry}} = 0.06 \text{ s}^{-1}$ ; $\Delta I_c = 25 \text{ mM}$<br>$k_{\text{ncov}} = 1.5 \text{ mM}^{-1} \text{ s}^{-1}$<br>$k_{\text{cov}} = 10 \text{ s}^{-1}$ | | $k_{\text{ncov}} = 1.5 \text{ mM}^{-1} \text{ s}^{-1}$ (estimated)<br>$k_{\text{cov}} > 8000 \text{ s}^{-1}$ | |
| $I_{\text{out}}$ | 7 ms | Fig. 7 b blue dotted line | 7 ms | Fig. 7 a blue line |
| $I_{\text{in}}$ | 20.1 ms | Fig. 7 b blue line | not observed | |
| E:I formation | 21.2 ms | Fig. 7 b orange line | not observed |  |
| E-I* formation | 93.4 ms | Fig. 7 b green line | 11.5 ms | Fig. 7 a green line |
| <sup>a</sup>from Eqns. 4, 5 and 6. |  |  |  |  |

**Figure 7.**
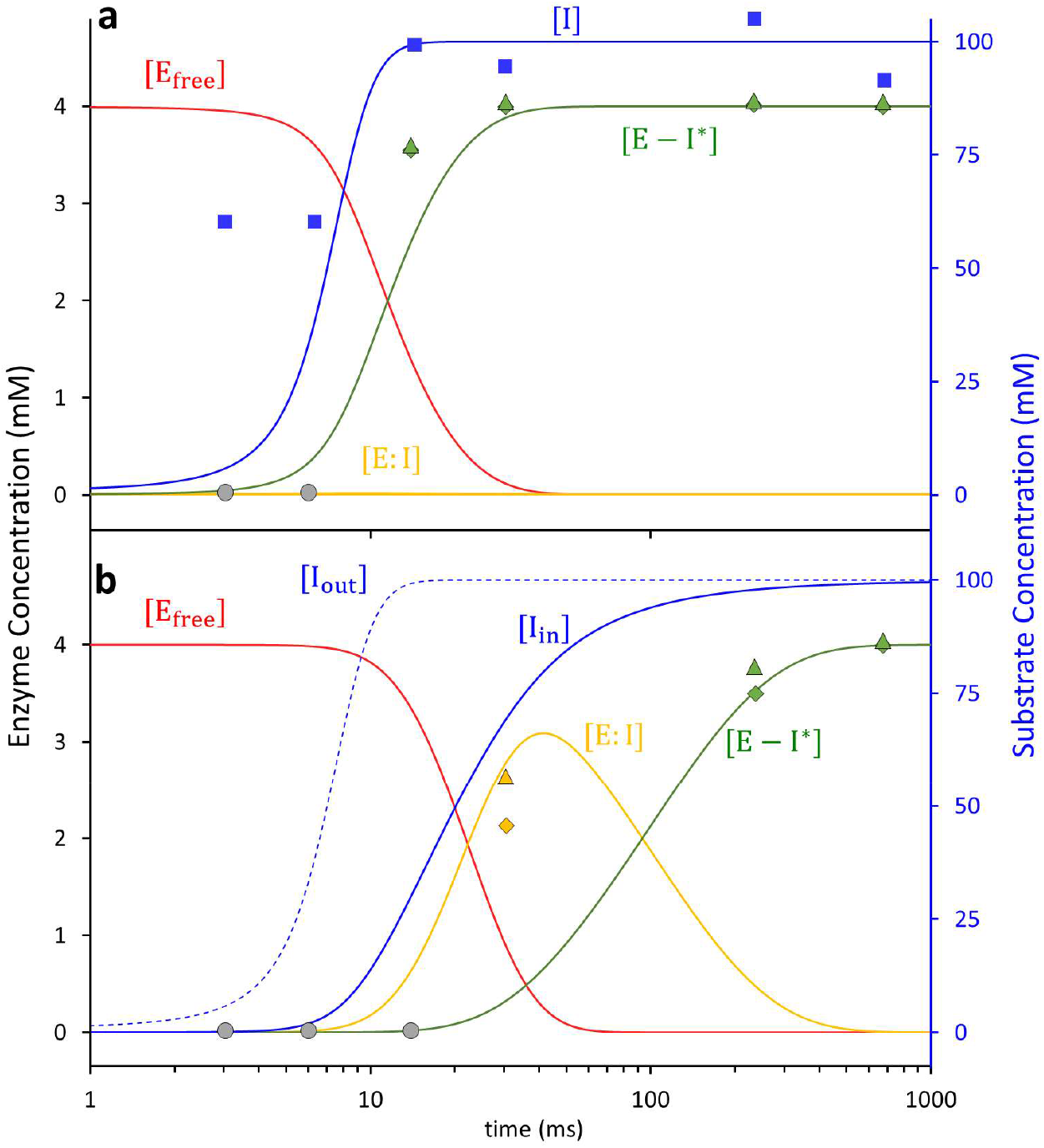
Calculated concentration profiles of reactants and products in the active sites of BlaC compared to corresponding observables. (a) Subunits B/D. Blue line: free SUB concentrations [I] in the unit cell. Blue squares: SUB concentrations in the central flow of the injector. Red line: time-dependent concentrations of the free BlaC [E_free_]. Orange line: concentrations of the non-covalently bound SUB [E:I] intermediate (not observable). Green line: concentrations of the covalent enzyme-inhibitor complex TEN [E-I^*^]. Green triangles and diamonds: concentrations of E-I^*^, derived from refined ligand occupancy values in subunits B and D, respectively. Grey circles: SUB cannot be detected near the active sites. (b) Subunits A/C. Blue dotted line: free SUB concentrations [I] in the unit cell. Blue line: SUB concentrations [I_in_] in the active site (note the delay relative to subunits B/D). Red line: time-dependent concentration of the free BlaC [E_free_]. Orange line: concentrations of the non-covalently bound SUB [E:I] intermediate. Green line: concentrations of the covalent enzyme-inhibitor complex TEN [E-I^*^]. Orange and green triangles and diamonds: concentrations of E:I and E-I^*^ derived from refined ligand occupancy values in subunits B and D, respectively.

The concentrations of the free SUB within the microcrystals rise slightly slower than the values estimated in the central flow of the injector (compare Fig. 7 a blue solid line and blue squares). The use of a logistic function (Eqn. 3) that describes the ligand increase in the unit cell is justified in particular by (i) the steep first phase observed in the rSVs (Fig. 6), but also by (ii) the very rapid increase of the ligand in the inner flow of the injector constriction where saturation occurs already after 15 ms. The covalently bound TEN accumulates rapidly with a characteristic time of 11.5 ms (Tab. 4). No additional intermediate is observed.

In subunits A/C, the apparent diffusion time of SUB necessary to reproduce the occupancies of the non-covalently bound intermediate is 20 ms (Fig. 7 b and Tab. 4). This is much longer than that observed in subunits B/D (7 ms). This lag can only be explained by a restricted access to the active site. The guardian residues open the active site after about 6 ms which corresponds to about 35 mM outside ligand concentration (Fig. 7 a). Entry to active site is controlled by an additional rate coefficient, k_entry_. k_entry_ was modelled (Eqn. 5) by an exponential function that depends on the (time-dependent) concentration difference Δ*I* (*t*) = *I*_*out*_ (*t*) − *I*_*in*_ (*t*) between the outside and the inside of the active site, respectively, and a characteristic concentration difference ΔI_c_ set to 25 mM (Tab. 4). I_out_ is the SUB concentration in the unit cell, which is known from the substrate binding kinetics to subunits B/D (see above). After a delay of ∼20 ms, SUB enters the active sites of subunits A/C and I_in_ rapidly increases. On the same time-scale the non-covalent E:I intermediate accumulates (Fig. 7 b). The non-covalently bound SUB triggers the next step of the reaction which results in a covalently bound TEN. The concentration of TEN ([E-I^*^]) starts dominating [E:I] at around 90 ms. By 240 ms, more than 90 % of the crystal is occupied by TEN (Fig. 7 b).

## 5. Discussion

### 5.1 Sulbactam forms a *trans-*enamine complex with BlaC

The acyl moieties are known to tautomerize into more stable *cis*- or *trans*-enamine products. In accordance, the SUB bifurcates into *cis-* and *trans-*enamines caused by the isomerization about the C5=C6 double bond ^55,56^ (Fig. 1 c, d). The presence of both products has been reported from spectroscopic data favoring the *trans*-enamine form^41,57^. It is argued that the further isomerization of the *cis*- to the *trans*-form is promoted by intrinsic steric clashes of the *cis-*form ^22,58^. At the resolution achieved here (between 2.35 Å and 3.2 Å, Tab.1), it is important to correctly discriminate between these two forms. The carboxylic and sulfinic moieties branch out from the common stem but they would form a similar shaped electron density in either of the conformations. To address this concern, a *cis-*enamine was placed in the DED_omit_ maps and refined (Fig. 8). The model poorly fitted the DED, and worse R-factors were obtained (not shown). This shows that the *trans*-enamine is the only species that can explain the electron density properly.

**Figure 8.**
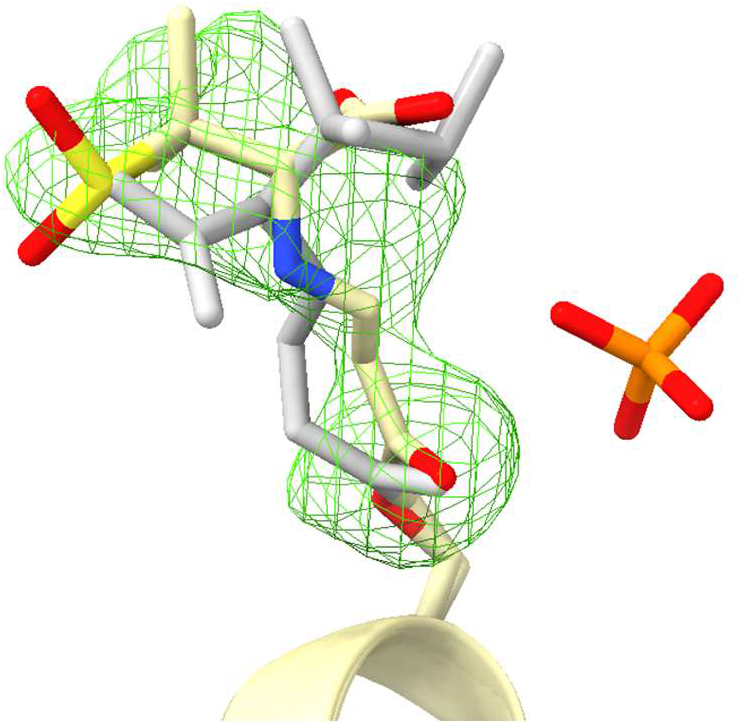
Stereo view of active site of subunit B. *Cis-*enamine (shown in grey) poorly fits the electron density at Δt_MISC_ = 15ms.

### 5.2 Comparison of kinetics in subunits B and D versus A and C

While it is established that all the subunits take part in the reaction despite the heterogeneity, they do so at different rates. In subunits A/C, BlaC is inhibited by SUB via a two-step mechanism. The non-covalent intermediate can accumulate since the SUB is not properly oriented (Fig. 3 d) and can react to TEN only after an additional time delay. SUB displacements will be restricted by interactions with the surrounding residues (Tab. 2). To react with the active Ser70, the SUB must re-orient to expose the β-lactam ring towards the serine. The catalytic opening of the β-lactam ring and the unfurling of the thiazolidine ring all happen in succession inside the narrow reaction center cavity. This severely lowers the rate of TEN formation.

In subunits B/D the covalent binding of SUB can apparently be explained by a one-step mechanism. However, the two-step mechanism described is also consistent with the observations when the non-covalent binding of substrate to the enzyme is much slower than the formation of TEN. By applying the two-step mechanism (Eqn. 4), a k_ncov_ value of ∼1.5 mM^-1^ s^-1^ and a large k_cov_ value of ∼8000 s^-1^ are estimated (Tab. 4) so that the non-covalently bound E:I complex does not accumulate. The open binding pocket accommodates for large chemical and structural changes which result in the large k_cov_. The rate determining step is controlled by k_ncov_ although the non-covalent BlaC-SUB complex never accumulates. In a one-step scenario (the free SUB reacts directly to TEN), the pseudo one-step rate coefficient would have to be the same as the k_ncov_, which was confirmed with a separate calculation (not shown). The two-step mechanism explains the enzymology in the active sites of all subunits in a consistent way.

A second relaxation phase is observed in subunits B/D, although the reaction to TEN has already taken place. It appears as if this is a result of the continuing reaction in subunits A/C and, related to this, an ongoing relaxation of the entire protein structure. The relaxation times, τ_2_ of the second relaxation phase are all between 80 ms and 90 ms for the four subunits (Tab. 3). The relaxation times are tied together, which is an indication for a cooperative behavior of all four subunits. This observation would have been obscured without the prowess of SVD which can track DED values across the entire reaction with great precision.

### 5.3 Rapid diffusion with Sulbactam in BlaC microcrystals

By taking into account the molecular volume that is enclosed by the van der Waal’s surface, SUB (181.7 Å^3^) is 2.5 times smaller than CEF (444.9 Å^3^). For CEF, 50% occupancy of the enzyme substrate complex has been observed at Δt_MISC_ = 5 ms in subunits B and D ^34^. Due to the smaller size, SUB should diffuse faster into the crystals than CEF and the signature of an enzyme inhibitor complex is expected to appear already at the earliest time points (3 ms and 6 ms). However, the first event that could be identified in the DED maps is the appearance of the TEN at 15 ms in subunits B/D (Fig. 3 i). The concentrations of SUB at the active site of subunits B/D, which allow direct, unrestricted access, can be taken as an estimate of the SUB concentration inside the BlaC crystals. The slight lag between the estimated SUB concentration in the inner flow of the injector constriction and the concentrations in the crystals at early time points (Fig. 7 a, compare blue squares with the blue line) is expected, since diffusion in BlaC crystals is slowed down in crystals ^34^ compared to water. The apparent diffusion time (7ms, Tab. 4) of the SUB is almost the same as that for the antibiotic substrate CEF (∼ 5 ms)^34^. Still, contrary to expectations based on the size of the SUB, no electron density has been present at the earliest time points. This can be explained by the smaller second order binding coefficient, k_ncov_ (1.5 mM^-1^ s^-1^ for SUB compared to 3.2 mM^-1^ s^-1^ for CEF), that prevents the accumulation of electron density at earlier times.

The characteristic times observed in both classes of subunits for the formation of the covalently bond inhibitor species (around 10 ms and 90 ms, respectively) are quite fast in comparison to earlier works that were suggesting that the reaction might take minutes to complete ^27,41^. The fast reaction provides an advantage when β-lactam substrates and the SUB inhibitor are competing for the same active site. For example, the non-covalent enzyme-substrate complex with CEF persists for up to 500 ms ^33^. During this time, the CEF can leave the enzyme and be replaced first competitively and then irreversibly by a quickly reacting SUB molecule. Since the inhibitor competes with co-administered antibiotics for the active site of BlaC, one can imagine that the covalent bond formation with an inhibitor must occur as fast as possible to effectively eliminate β-lactamase activity in the presence of substrate. This is in addition corroborated by Jones and coworkers who reported that SUB has a ten times higher affinity and binding constant for plasmid mediated class A β-lactamases compared to cefoperazone ^59^, which is a third-generation cephalosporin-based antibiotic similar to CEF.

### 5.4 Ligand gating, induced fit and conformational selection

Our results show that ligand binding to enzymes may be more complicated than initially thought ^27,37,38^. Only after a delay the ligand penetrates into the active sites of subunits A/C controlled by the guarding residues (Fig. 7 b). The narrow entrance to active site (Fig. 2 b), and the displacements of the guardian residues (Tab. 2) are reminiscent of a substrate tunneling-and-gating ^7-9^ like mechanism which has not yet been discovered in published structures of BlaC. More work is needed to determine the mechanism that drives the displacement of these residues. As these displacements were not observed when reacting with CEF ^33,34^, an allosteric mechanism that links the position of the guardian residues to the covalent binding of SUB in adjacent subunits is unlikely. However, electrostatic interactions ^60^ of the negatively charged sulbactam with the positively charged Arg173 or even polar interaction with the Gln112^B,D^, respectively, may plausibly induce these structural changes.

At Δt_MISC_ of 30 ms and 66 ms ^34^ SUB occupies the active sites of subunits A/C without reacting with Ser70. During this time conformational changes of the BlaC are apparent (Tab. 2) to accommodate the SUB. This resembles an induced fit ^10^. The unfavorable SUB orientation prevents the direct attack of the active Ser70 towards the β-lactam ring. Ser70 can react with the β-lactam ring only after a rotation of the SUB. Fluctuations of the Ser70 towards the SUB carbon C_7_ will then lead to a very short lived, indiscernible “transition state like” structure E-I with a covalent bond between the SUB and BlaC. This is a different form of conformational selection ^12,13^, in a sense that there is not a particular “preferred” protein conformation that reacts with a substrate, but here, a particular (active) ligand orientation is required and “selected” by the enzyme for further reaction. It is the rate of the reorientation (rotation of the SUB) that seems to control the rate of this reaction. Once a favorable orientation is reached, further reaction to TEN is instantaneous on the timescale of the observation (90 ms) (Tab. 4). This informs the design of improved (faster) inhibitors that consist of symmetric active moieties ^61,62^ or are engineered to enter the active sites in the correct orientation.

In subunits B/D, the inhibitor is brought in rapidly by diffusion and reacts instantaneously (< 1.5 ms) on the timescale of observation (> 15 ms). Neither an induced fit nor conformational selection can be observed or distinguished which previously led to intense discussions for other enzymes ^11^. The active site structures relax in unison with the extent of covalently bound inhibitor in all subunits as explained above.

### 5.5 The fate of the *trans-*enamine

It has been proposed that on longer timescales (> 30 min) a second nucleophilic attack by a nearby serine can occur in other, structurally closely related Ambler Class A β-lactamases ^35^. This serine (Ser128 in BlaC) may react with the C5 position of the TEN (Fig. 1). This is followed by the loss of the opened thiazolidine ring fragment (Fig. 1 d). A covalent bond may be formed between C5 and Ser128 of BlaC (Fig. 9 a) leading to the prolonged inhibition of the enzyme ^35,41^. It has also been suggested that only the transient inhibition by TEN is responsible for the medical relevance of SUB as any reaction that lasts longer than one hour is irrelevant due to the bacterial lifecycle of ∼30 minutes ^41^. However, the life cycle of *Mtb* is around ∼20 hours ^63^. The permanent inhibition achieved only after the second nucleophilic attack might be the ultimate factor for SUB’s clinical usefulness in fighting antibiotic resistance in slow growing bacteria like *Mtb*. Inspection of the soaked structure may give an answer. Covalently bound TENs were observed in all four subunits when BlaC was soaked with SUB for 3 hours (Fig. 5). The B-factors of the fragment beyond N4 that would be cleaved off (displayed in pale colors in Fig. 9 a) are consistently higher by 20 Å^2^ than that of the part which would form the cross-linked species. However, it is more plausible that higher B-factors are caused by the dynamic disorder of the long TEN tail and not by the presence of mixture of intact and fragmented TEN. There is no clear evidence of TEN fragmentation, and TEN remains the physiologically important species for BlaC inhibition for hours.

**Figure 9.**
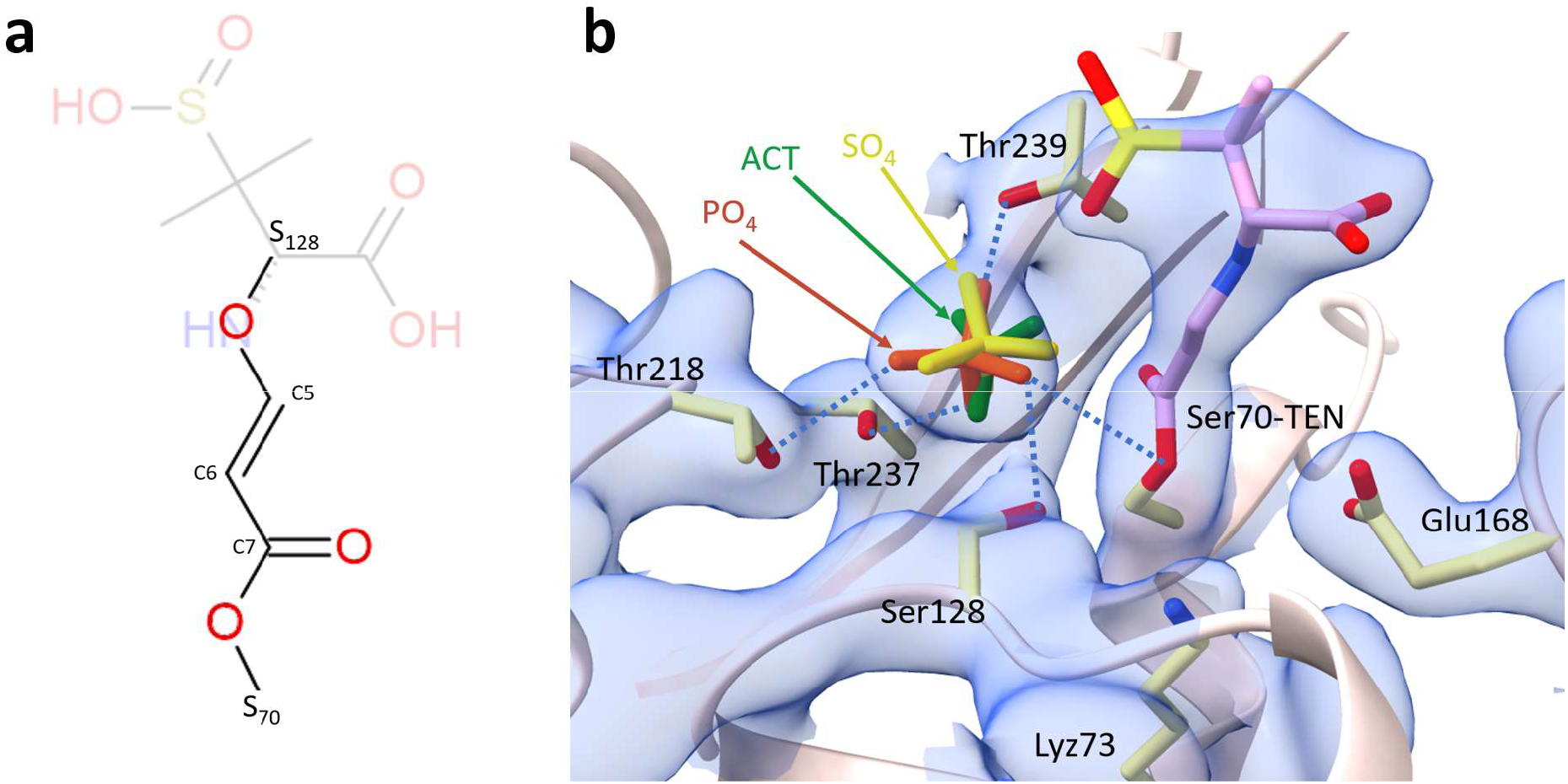
(a) Chemical structure of the TEN after the formation of cross-linked species. The leaving group is shown in pale color. (b) A 2F_obs_-F_calc_ map is shown near the active site of subunit B at Δt_MISC_ = 240ms (blue). The contour level is 1 σ. Key active site residues are marked. The phosphate molecule (PO_4_) is represented in orange. The hydrogen bonds made by phosphate with surrounding residues are depicted by blue dotted lines. PDB entries 5OYO and 7A71 are overlayed over the BlaC. An acetate (ACT, green) and a sulfate (SO_4_, yellow) occupy the same position as the PO_4_ in BlaC. TEN (purple) might not be able to get close to Ser128 without displacing the phosphate.

A phosphate group binds to a specific site immediately adjacent to the active serins 70 and 128 (Fig. 9 b) with multiple hydrogen bonds to surrounding residues. This location appears to be conserved among all published BlaC structures where others have also reported sulfate and acetate molecules in the same position (Fig. 9 b) ^27,64-66^. Naturally occurring compounds with phosphate like group, for instance adenosine phosphates, might also interact with BlaC. β-lactamase production increases in some bacteria grown in a phosphate enriched medium ^67^ while phosphate can also promote the hydrolysis of the clavulanate inhibitor by BlaC ^66^. More structures are required after soaking with high SUB concentrations for longer periods of time, perhaps days, to observe potentially fragmented TEN. Since the phosphate is replaceable ^1,33,34^, the TEN might indeed react further. Larger inhibitors like tazobactam and clavulanic acid might also be able to displace the phosphate molecule directly. Recently, *trans-*enamine intermediates of tazobactam were identified in the serine β-lactamase TEM-171 at a position similar to that of TEN in BlaC and at another that is occupied by the phosphate in BlaC ^68^. The diverse chemistry that is already observed very early on in BlaC may extend to much longer time scales.

### Other β-Lactamases

There are other classes of β-lactamases that are more concerning than BlaC such as the metallo β-lactamases (MBL). They are capable of hydrolyzing almost all clinically available β-lactam antibiotics and inhibitors ^69,70^. Similar work to the one presented here and earlier ^33,34^ could be performed on MBLs to gain more structural insight into their catalytic mechanisms. Time-resolved pump-probe crystallographic experiments using a caged Zn molecule ^71^ already show how the antibiotic moxalactam is inactivated by a MBL on a timescale longer than 20 ms. To characterize the important substrate binding phase on single ms, and even sub-ms time scales, it would be desirable to follow this or a similar reaction as well as the binding of a MBL inhibitor with MISC.

## 6. Conclusion and outlook

MISC is a straightforward way to structurally study enzyme function. Reactions are visualized in real time as movies of the enzyme in action. Our results give insight how the shape of the active site determines rate coefficients and reaction mechanisms of a biomedically relevant reaction. From the MISC data, diffusion times and rate coefficients can be estimated by applying informed, chemically meaningful constraints and tying the analysis to observed occupancy variations of transient enzyme-inhibitor complexes in the different subunits. The key advantage of MISC is that it provides at the same time a localized view onto processes that unfold in individual subunits and a global perspective of the behavior of the entire molecule with near atomic precision. High repetition rate XFELs ^72,73^ and upgraded synchrotron light sources ^74,75^, will facilitate the collection of time-series that consists of very closely spaced MISC time-delays. A global evaluation of these time-series assisted by a principal component analysis such as the SVD and other machine learning techniques ^47,49^ will provide a detailed and direct view into enzyme catalysis and inhibition.

## 7. Methods

### 7.1 Sample preparation

BlaC was expressed in E. coli and purified as previously described ^33^. Crystals were grown on site at the Linac Coherent Light Source (LCLS) as published ^33^. In short, purified BlaC protein (concentrated to 150 mg/ml) was added dropwise to 2.4 M ammonium phosphate (AP) at pH 4.1 while stirring until a ratio of 1:9 (protein: precipitant) was reached. The stirring was stopped after ∼12 hours. Protein microcrystals were left to mature for 2 days at room temperature. As the crystals settled, the supernatant was removed to reach the desired concentration. To avoid potential clogging of the injector nozzles, the crystal slurry was filtered through a 20-micron syringe filter. The final concentration of crystals loaded into the injector reservoir was 5 × 10^9^ crystals/ml. For mixing, a solution of 150 mM SUB in 50mM AP at pH 4.5 was prepared. A reference (unmixed) dataset was obtained by mixing the BlaC microcrystals with water.

### 7.2 Data collection and processing

Data were collected at the Macromolecular Femtosecond Crystallography (MFX) instrument ^76,77^ at the LCLS in October 2020 [beamtime lu6818]. The XFEL was operating at 120 Hz with a pulse energy of ∼9.8 keV (λ = 1.26 Å) and a pulse duration of 20-40 fs. The XFEL beam was focused to a spot size (FWHM) of 3μm. The crystal slurry was mixed with SUB solution and injected into a helium filled chamber at ambient temperature. The injectors allow simultaneous mixing and serial injection of sample into the X-ray interception region. Injectors were provided by Pollack group (Cornell University) and are based on the design published by Calvey et al. ^78,79^. A central thin stream of a suspension containing microcrystals flows down a delay line of variable length, called the constriction (Fig. 10). A solution of substrate or ligand concentrically surrounds the central stream allowing for rapid diffusion into the microcrystals within the stream^79^. At the end of the constriction, the mixture is injected into the X-ray interaction region. The schematic and geometry of the injectors are shown in Fig. 10 and Tab. 5 respectively.

**Figure 10.**
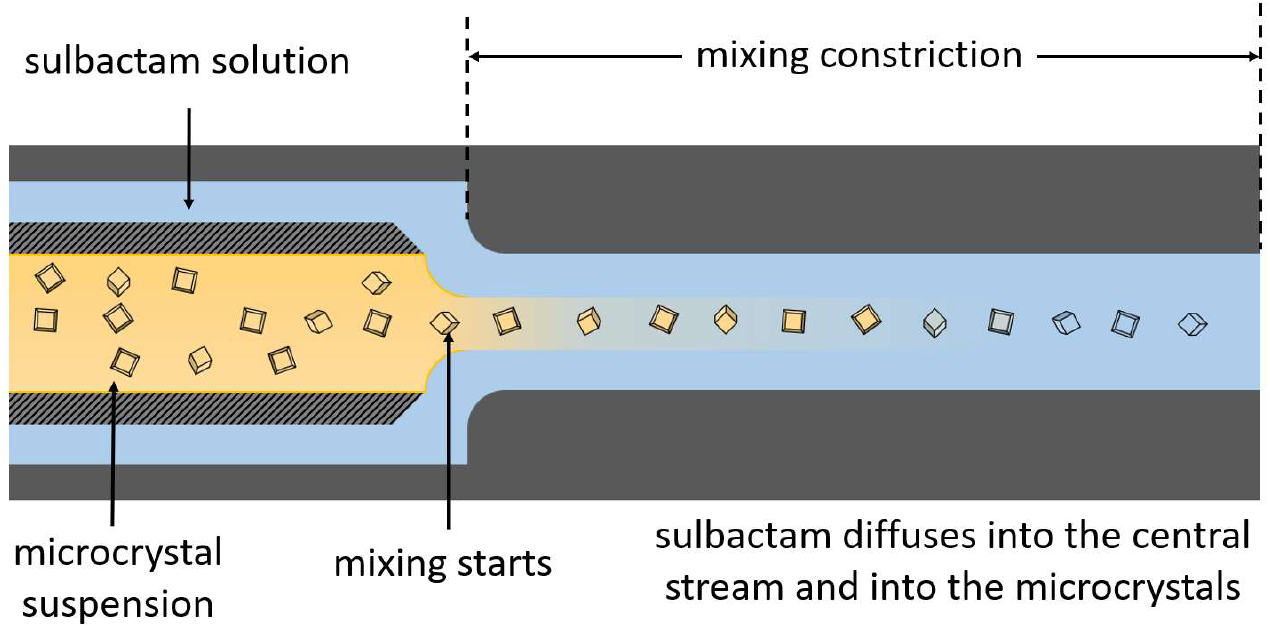
Cartoon cross section of the injector focused on the mixing region adapted from Calvey et al.^40^. Randomly oriented BlaC microcrystals flow through an inner capillary, whereas the sulbactam solution flows concentrically through the outer capillary. Both flows are combined into a single outlet of reduced diameter called the mixing constriction. The time taken by crystals to move from the start of the mixing constriction until they are injected into the X-ray interaction region determines the MISC time delay. The sulbactam first diffuses into the central flow containing crystals, and then into the crystals. ∼100 mM of ligand concentration in the central flow is reached after 15 ms.

**Table 5.** Injector geometry and sample flow rates

|  | Water | Sulbactam |  |  |  |  |  |
| --- | --- | --- | --- | --- | --- | --- | --- |
| $\Delta t_{\text{MISC}}$ (ms) | 0 | 3 | 6 | 15 | 30 | 240 | 700 |
| Ligand concentration (mM) | n.a. | 150 |  |  |  |  |  |
| Ligand buffer | water | 50 mM Ammonium Phosphate (pH 4.5) |  |  |  |  |  |
| Injector ID | 3 | 1 | 1 | 2 | 2 | 3 | 4 |
| Crystal flow rate ($\mu\text{l}/\text{min}$) | 10 | 7.7 | 7.6 | 7.7 | 11.1 | 10.1 | 11 |
| Ligand flow rate ($\mu\text{l}/\text{min}$) | 70 | 142.3 | 67.4 | 143.1 | 63.1 | 70 | 57 |
| Constriction<sup>†</sup> inner diameter ($\mu\text{m}$) | 100 | 50 | 50 | 50 | 50 | 100 | 100 |
| Constriction length (mm) | 82.7 | 10.3 | 10.3 | 40.7 | 40.7 | 82.7 | 35<sup>‡</sup> |
| Timing uncertainty (ms) | n.a. | $\pm 1.0$ | $\pm 2.0$ | $\pm 1.1$ | $\pm 3.5$ | $\pm 18.8$ | $\pm 29.5$ |
| Ligand concentration at the end of constriction (mM) | n.a. | 58.5 | 58.5 | 100.5 | 88.5 | 112.5 | 99 |
| <sup>‡</sup>For timepoints longer than 500 ms, an additional delay stage is added immediately after constriction. |  |  |  |  |  |  |  |

Our BlaC crystals are quite small (∼10 × 10 × 2 µm^3^) and contain large solvent channels (Fig. 2 a), both of which promote rapid diffusion of substrate into the crystals. The time delay (Δt) in MISC is the time taken by the crystals to travel from the mixing region in the injector to the X-ray interaction region. “Time point” and “time delay” are used interchangeably throughout this manuscript and denoted by Δt_MISC_. Different injector designs allowed data acquisition at 6 different Δt_MISC_ from 3 ms to 700 ms (Tab. 1; Tab. 5).

Diffraction patterns (DPs) were collected with the epix10K2M detector ^80^ at 120 Hz. The experiment was monitored in real time with the OnDA (online data analysis) Monitor (OM) ^81^, which also provided feedback on hit rate and spatial resolution. The raw data were processed by Cheetah ^82^. Cheetah identifies DPs containing potential Bragg reflections (hits). In addition, it has a built-in masking tool that allows masking of unwanted detector pixels. The selected patterns were further processed by the CrystFEL suite of programs ^83,84^. *Indexamajig* was used to index and integrate the DPs using a combination of *mosflm, dirax, xds, asdf* and *xgandalf* ^85-88^. The detector geometry and distance were refined by *geoptimiser* ^*89*^. Merging and scaling of the intensities were performed with *partialator* using the partiality model *xsphere* ^90^. Figures of merit and other data statistics were calculated using *compare_hkl* and *check_hkl*.

Macroscopic crystals were grown in sitting drops (10µl BlaC at 45mg/ml^−1^ mixed (1:1) with 2.1 M AP at pH 4.1). Crystals were soaked for 3 hours in a cryobuffer consisting of 2 M AP, 20% glycerol and 100 mM SUB. The crystals were flash frozen in liquid nitrogen and investigated at beamline ID-19 of the Structural Biology Center, Advanced Photon Source, Argonne National Laboratory. Data were collected by the proprietary *sbccollect* program ^91^ and processed by *HKL-3000* ^92^. The data collection and refinement statistics are listed in Tab.1.

## 8. Data analysis

### 8.1 Difference maps and structure determination

Because of the change in unit cell parameters, omit difference electron density maps were calculated throughout as previously described ^33,34^. First, a reference (unmixed) structure was refined using the structure published in the protein data base (PDB) ^93^ entry 6B5X as initial model. For the analysis of the MISC data, the water and the phosphate molecules in the active sites were removed from the reference structure. For each Δt_MISC_, the resulting structure was refined using the standard simulated annealing (SA) protocol in *Phenix* ^94^ against the experimentally observed structure factor amplitudes |F_obs_(t)|. Following refinement, m|F_obs_(t)| -D|F_calc_| omit maps (DED_omit_) were calculated for each time point, where |F_calc_| are calculated from the refined model. Polder difference maps (PDMs) ^95^ were calculated to display weak ligand densities in some time points to assist in the placement of the ligand. To calculate a PDM, a suitable small molecule (which was SUB in our case) is placed at the center of the active site at a position where the ligand density is expected to be. The algorithm then generates an omit map by excluding bulk solvent modeling around the selected region. This way, weak densities become apparent, which may otherwise be obscured by the bulk solvent. PDMs were calculated particularly for the 30ms and the 700ms timepoint, due to the relatively low number of indexed DPs (Tab. 1), and for the cryosoaked data (Tab. 3).

Both the unbound sulbactam and the covalently bound *trans*-enamine (TEN) were manually modeled into the DED maps using *Coot* ^*96*^. The ccp4-program *AceDRG* ^97^ was used to generate coordinates and restraint for the covalently bound TEN. Several cycles of refinement in *Refmac* ^98^ and *Phenix* followed by manual inspection and re-modeling with *Coot* led to models with excellent R-factors. Ligand occupancies were determined by ‘group occupancy refinement’ in *Phenix*. A value of 100% indicates that stoichiometric concentration was reached. The refinement statistics are listed in Tab. 4. All figures that display DED maps were prepared using *UCSF ChimeraX* ^99^. The chemical structure drawing software *ChemSketch* (ACD/Labs) was used to create chemical schematics.

### 8.2 Singular value decomposition

A kinetic analysis was performed by the application of the singular value decomposition (SVD) to the X-ray data ^14^ using the DED_omit_ maps. A region of interest (ROI) was determined that corresponds roughly to the volume of an individual active site selected from the four subunits in the asymmetric unit. The ROI covers the volume occupied by the SUB and TEN ligands and the entire side chain of Ser70 (CA, CB and OG). It touches the terminal atoms of residues Lys 73, Glu168, Thr239, Asn172, Arg173, Ser128, Ser102 and Gln112 (from B/D) in the case of subunits A/C. Difference electron density values in the region of interest were assigned to a m-dimensional vector. N of these vectors were obtained for all measured Δt_MISC_ and assembled to a m x n dimensional matrix **A**, called the data matrix. SVD is the factorization of matrix **A** into three matrices: **U, S** and **V**^**T**^ according to

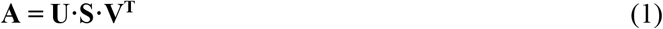

The columns of the m × n matrix **U**, are called the left singular vectors (lSVs). They represent the basis (eigen) vectors of the original data in data matrix **A. S** is an n × n diagonal matrix, whose diagonal elements are called the singular values (SVs) of **A**. These non-negative values indicate how important or significant the columns of **U** are. The columns of n × n matrix **V**, called the right singular vectors (rSVs), contain the associated temporal variation of the singular vectors in **U. S** contains n singular values in descending order of magnitude.

The number of significant singular values can inform how many kinetic processes can be resolved. The SVD results can then be interpreted by globally fitting suitable functions to the rSVs, that can consist, in the simplest form, of sums of exponentials. The rSVs contain information on the population dynamics of the species involved in the mechanism ^14,100^.

The earlier program SVD4TX ^14^ and a newer version ^101^ could not be applied to X-ray data when large unit cell changes occur during the reaction since these implementations relied on a region of interest that is spatially fixed. This is not given when the unit cell changes. In order to accommodate changing unit cells, a new approach was coded by a combination of custom bash scripts and python programs described below.

### 8.3 Adapting SVD for MISC Datasets with Changing Unit Cell Parameters

The DED map are calculated in the CCP4 file format ^102^ and cover the entire unit cell of the crystal. The maps are represented by a three-dimensional (3D) array with m_x_, m_y_ and m_z_ grid points for each unit cell axis, respectively. Each 3D grid point (voxel) contains the value of difference electron density at that given position (Fig. 11 a). In such a DED map, the green (positive) features indicate regions where atoms have shifted away from their position in the reference model. Negative (red) features are then found on top of the atoms in the reference model. Most of the map contains spurious noise except in ROIs such as the active sites where larger structural changes are expected due to the binding or dissociation of a ligand (Fig. 11 b). The noise within the majority of the difference map would interfere with the SVD analysis. To avoid this, a ROI was isolated individually for each subunit and an SVD performed only on the DED within. When multiple active sites are present, each active site can be investigated separately. Fig.12 shows a flow chart of the steps required to prepare the data matrix **A**. The steps are described in detail below.

**Figure 11.**
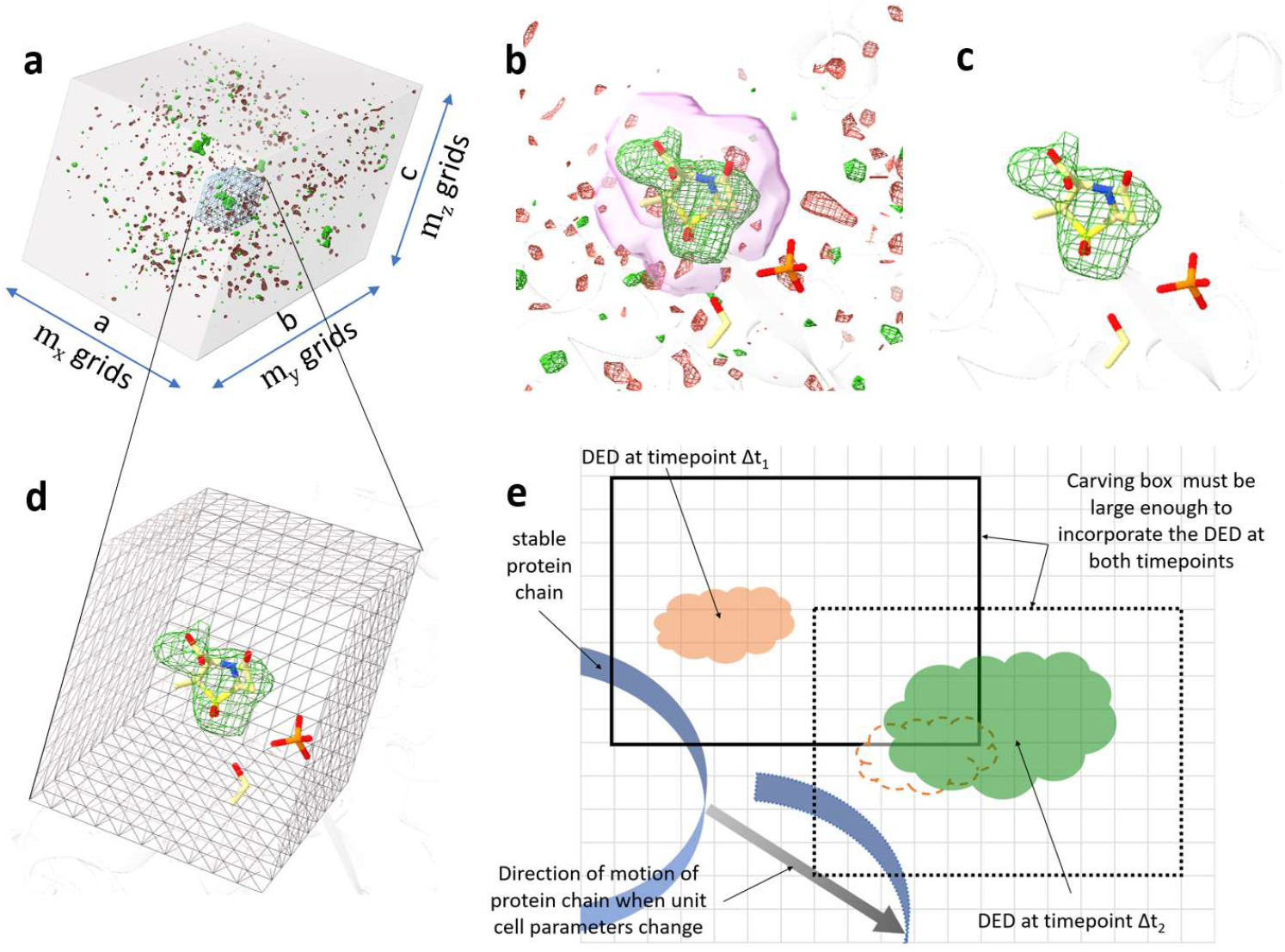
Carving out the region of interest (ROI) from DED maps when unit cell parameters change. (a) DED map of the entire BlaC unit cell at Δt_MISC_=30ms contoured at ±2.5 σ (b) The same map as in (a) is now displayed with the focus on the active site of subunit A. Strong DED is present where the SUB molecule resides. A mask around the SUB atom is represented by a pink surface. The density inside the mask is left unaltered while that outside the mask is set to zero. (c) The map after the masking operation. (d) A box which is a part of overall map covers the ROI. (e) A simple 2D diagram showing how to choose the box. The square grey grids represent the voxels. The box shown by the solid line includes the ROI at Δt_1_ and it is large enough to cover the evolving DED features at all other time points. The orange cloud represents the DED features at timepoint Δt_1_. At Δt_2_, the entire protein chain displaces to a new position (grey arrow) due to unit cell changes. In addition, more extensive DED features appear as shown by the green cloud. The dashed cloud next to the green DED is the relative position of the DED at Δt_1_. As the protein chain displaces when the unit cell parameters change, the entire box moves accordingly in the same direction (shown by the dotted box). Since the number of grid points are adjusted linearly with the changing unit cell parameters, the number of voxels within the box as well as the voxel size does not change.

**Figure 12.**
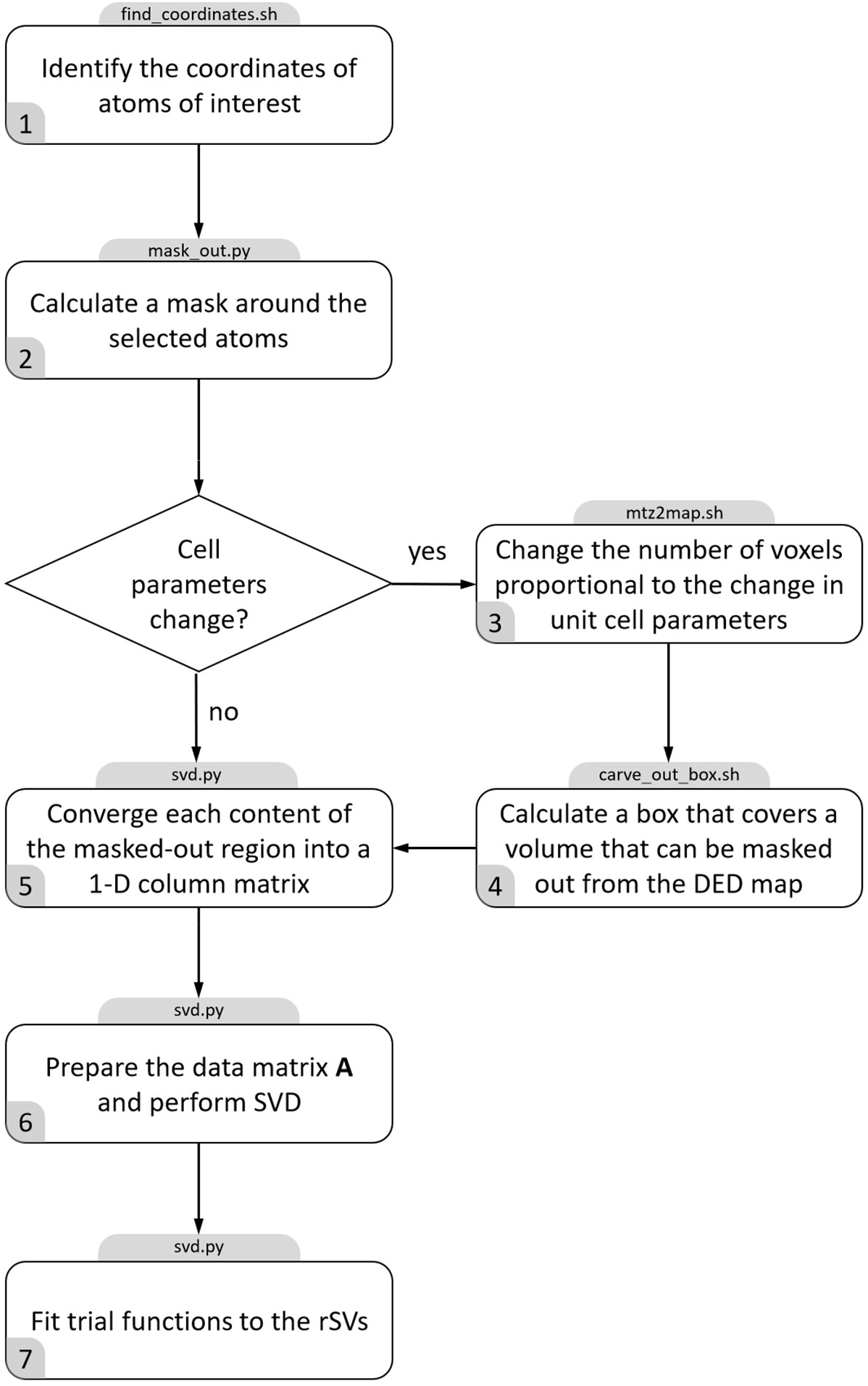
Flow chart for an SVD analysis of time-dependent crystallographic difference maps. The grey boxes on top are the scripts/programs developed and used to accomplish each step.

**Step 1:** The coordinates of the atoms of the amino acid residues and the substrate of interest are specified in a particular subunit. This defines the ROI. For the present work, four different ROIs were defined, one for each subunit A to D, respectively, and investigated separately.

**Step 2:** A mask is calculated that covers the selected atoms plus a buffer distance of choice (Fig. 11 b). The density values outside of the mask are set to 0 while the ones inside are left unchanged. This results in a masked map with the dimensions of original map with density values present only around the emerging DED in the active site (Fig. 11 c). This mask was evolved later (after step 4) by allowing only grid points that contain DED features greater or smaller than a certain sigma value (for example, plus or minus 3 σ) found at least in one time point ^14^.

When the unit cell parameters do not change during the reaction, the difference maps at all time points will have the same number of voxels and the voxel size is also constant. However, once the unit cell dimensions change, either the voxel size will change, if the number of voxels is kept constant, or the number of voxels will change, if the voxel size is kept constant. If the voxel size changes, the DED value assigned to each voxel position will also change which will skew the SVD analysis. If the voxel numbers change, the SVD algorithm will fail as it requires that all the maps are represented by arrays of identical sizes. Accordingly, both conditions, (i) a constant voxel size and (ii) a constant number of grid points in the masked volume must be fulfilled when the unit cell changes.

**Step 3:** In order to fulfill (i) the total number of grid points in the DED map is changed proportionally to the unit cell change. When the volume of the ROI is not changed, condition (ii) is automatically fulfilled, and a suitable data matrix **A** can be constructed. However, when the unit cell parameters change, the ROI is also changing position. This must be addressed in addition.

**Step 4:** A box is chosen that will cover the density that was just masked out (Fig. 11 d). The box will include the ROI to carve out the DED values which is saved as a new map. The size of the box must be large enough such that the ROIs can be covered at all time points. The box must be calculated with reference to a stable structure (usually the protein main chain). As the protein chain displaces as a result of the change of the unit cell, the box will also move accordingly to cover the correct ROI (Fig. 11 e). As mentioned, the DED within the moving box can be used to evolve the mask that defines the final ROI as indicated in step 2.

**Step 5:** All m voxels in the evolved mask are converted to a one-dimensional (1D) column array, a vector in high (m) dimensional space. How the conversion is achieved does not matter as long as the same convention is applied to all the n maps. The reversion of the protocol can reconstruct a 3D difference map (for a particular subunit) from the m-dimensional

527 vector. N of the m-dimensional vectors are arranged in ascending order of time to construct the data matrix **A**.

**Step 6:** SVD is performed on matrix **A** according to Eqn. 1.

**Step 7:** Trial functions are globally fit to the significant rSVs to determine relaxation times and the minimum number of intermediates involved in the reaction (see e.g., Ihee et al., 2005 ^53^).

### 8.4 Global fit of the Significant rSVs

For a simple chemical kinetic mechanism with only first-order reactions, relaxations are characterized by simple exponentials ^48^. For higher order reactions, the rSVs have to be fitted by suitable functions which must explain the changes of the electron density values in a chemically sensible way ^14,29,103^. In our case, the significant rSVs were fitted by the function shown in Eqn. 2 which, apart from a constant term, consists of a logistic function that accounts for abrupt changes of the electron densities observed in the active sites, and a saturation term. Further, the fit was weighted by the square of the corresponding singular values S_i_.

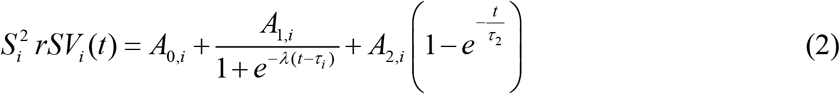

While the amplitudes A’s are varied independently for each significant rSV_i_ (i = 1…n), the parameter λ and the relaxation times, τ_1_ and τ_2_ are shared globally. The number of relaxation times (here 2) is equal to the number of significant rSVs and to the number of distinguishable processes.

### 8.5 Species Concentrations

The diffusion of SUB molecules into the BlaC crystals and subsequently into the active sites triggers the reaction. Concentrations of SUB in the central flow were estimated according to Calvey et al., 2019 ^79^. At the longest Δt_MISC_= 700 ms, the SUB concentration was 100 mM, which was used as the maximum ligand concentration for all calculations. The resulting evolution of concentration of SUB at the active sites was modeled by Eqn. 3.

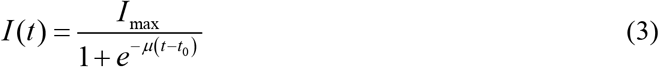

I(t) is the concentration of SUB at the active sites averaged over all unit cells in the crystal as a function of time and I_max_ is the maximum (100 mM) SUB concentration (Tab. 5). Eqn. 3 is a logistic function where μ is the growth rate and t_0_ is the midpoint value of the growth.

Once the SUB molecule reaches the active site of BlaC, the first step is the formation of a non-covalent enzyme inhibitor complex (E:I) (Fig. 13). The process depends on the free BlaC concentration inside the crystal, and the rate coefficient for non-covalent complex formation (k_ncov_). This step is usually reversible defined by both the forward rate coefficient (k_1_) and the backward rate coefficient (k_-1_). However, the mounting concentrations of inhibitor inside the crystals forces more molecules towards the active site. At least initially, the binding rate depends on k_1_ alone. The non-covalent E:I complex is the reactant for the next phase of the reaction where the β-lactam ring opens. The resulting covalently bound acyl-enzyme complex (E-I) (Fig. 13) is so short lived that it never accumulates in the timescale of the measurement. The SUB undergoes rapid modification, and a product is formed where the enzyme is covalently bound to the irreversibly modified inhibitor (E-I^*^). k_cov_ is the apparent rate coefficient which describes the velocity of E-I^*^ formation directly from the E:I complex (Fig. 13). Ligand concentrations were determined by numerically integrating the following rate equations that describe the mechanism in Fig. 13.

**Figure 13.**
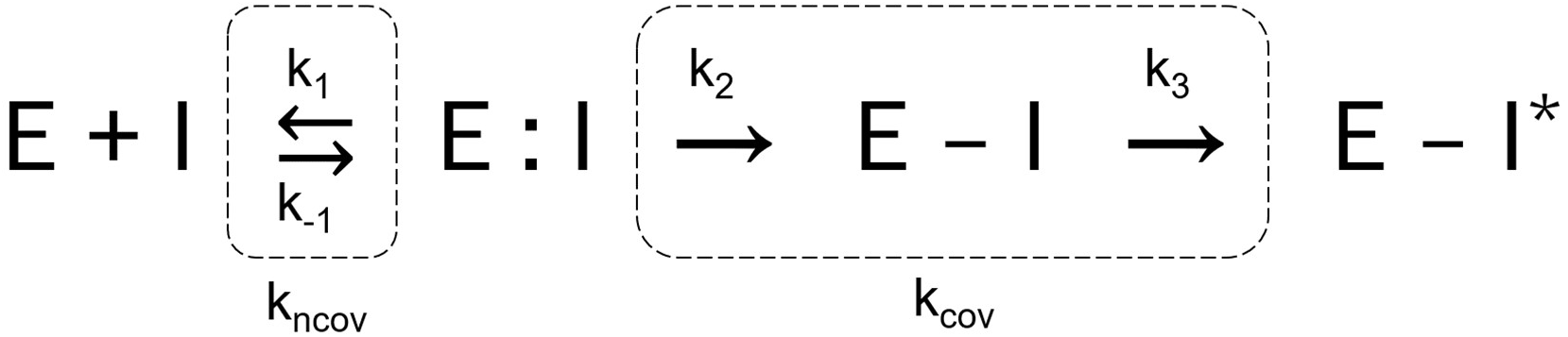
A simplified two-step mechanism of BlaC inhibition by sulbactam. The first step is the formation of the non-covalent enzyme inhibitor complex whose rate of formation depends on the concentration of the inhibitor in the unit cell and the rate coefficient (k_ncov_). The nucleophilic attack by the active serine opens the lactam ring of SUB leading to the formation of acyl-enzyme intermediate (E-I). The E-I intermediate does not accumulate to become observable. The next step is the irreversible inhibition of enzyme by the chemically modified inhibitor (E-I^*^) which depends on the apparent rate coefficient (k_cov_).nsi

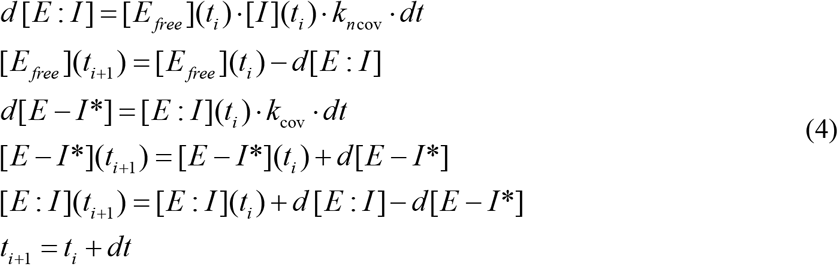

d[E:I] is the change in concentration of the non-covalent BlaC-SUB complex at any given time (t_i_) and depends on the free enzyme concentration, [E_free_], the second order rate coefficient of non-covalent binding (k_ncov_), and the inhibitor concentration, [I]. [I] is calculated from Eqn. 3. As the concentration of [E:I] increases, [E_free_] decreases. d[E-I^*^] is the increase of the covalently bound TEN. It depends on the available concentration of the non-covalent BlaC-SUB complex [E:I], and the rate coefficient k_cov_. [E:I] decreases by the same rate [E-I^*^] increases.

The increase of the SUB concentration in the active site (I_in_) is delayed relative to that of the SUB concentration in the unit cell (I_out_). To account for this delay, the rate coefficient that determines the entry into the active site (k_entry_, Fig. 14 a) is assumed to be dependent on the concentration difference

**Figure 14.**
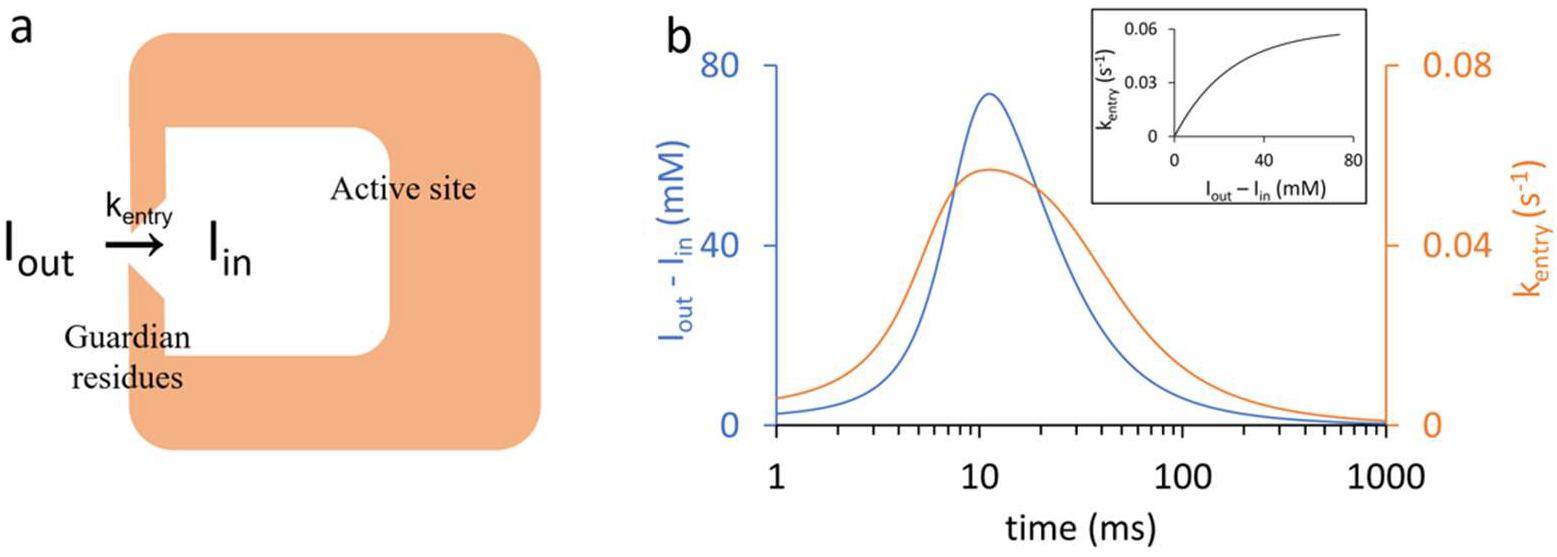
(a) Simplified scheme depicting the delayed entry of sulbactam into the active site through the guardian residues. (b) Time-dependence of the concentration difference (blue line) and of the rate coefficient k_entry_ (orange line). At early MISC delays k_entry_ is small. The channel opens, and k_entry_ is large only when sufficient SUB has accumulated in the unit cell Inset: The dependence of k_entry_ on the concentration difference.

Δ*I* (*t*) = *I*_*out*_ (*t*) − *I*_*in*_ (*t*) between outside and inside the active site and a characteristic difference Δ*I*_*c*_. It is modeled by an exponential function as shown.

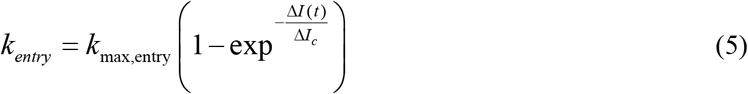

The relevant SUB concentrations within the active site (I_in_) are generated by solving the following rate equation:

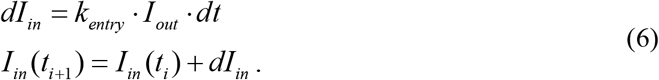

Eqn. 6 is used in lieu of Eqn.3 to calculate the relevant inhibitor concentration: I_in_(t) is fed as [I](t) to Eqn. 4 to calculate the concentrations of the non-covalently and covalently bound species shown in Fig. 5 b. At early MISC delays k_entry_ is small. The channel opens, and k_entry_ is large only when sufficient SUB has accumulated in the unit cell (Fig. 14 b). All relevant parameters are listed in Tab. 4.

## Acknowledgement

This work was supported by NSF-STC-1231306 (BioXFEL). P.F. was supported by NSF BioXFEL STC grant NSF-1231306 Biology with X-Ray Lasers the NIH grant R01GM095583 and the ASU Biodesign Center for Applied Structural Discovery. A.O. was supported by the US Department of Energy, Office of Science, Basic Energy Sciences under award DE-SC0002164 (underlying dynamical techniques), and by the US National Science Foundation under awards STC-1231306 (underlying data analytical techniques) and DBI-2029533 (underlying analytical models). K.A.Z. was supported by the Cornell Molecular Biophysics Training Program (NIH T32-GM008267). We acknowledge funding from DESY (Hamburg, Germany), a member of the Helmholtz Association HGF; the Cluster of Excellence “Advanced Imaging of Matter” of the Deutsche Forschungsgemeinschaft (DFG) - EXC 2056 - project ID 390715994; the Helmholtz Association Impulse and Networking fund - project InternLabs-0011 “HIR3X”; the German Federal Ministry of Education and Research (BMBF) - project 05K18CHA. Use of the LCLS, SLAC National Accelerator Laboratory, is supported by the U.S. DOE, Office of Science, BES, under contract no. DE-AC02-76SF00515. The HERA system for in helium experiments at MFX was developed by Bruce Doak and funded by the Max-Planck Institute for Medical Research. The structure factors and the refined coordinates of XFEL structure of BlaC mixed with sulbactam for 15ms, 30 ms, and 240ms, and the cryo-soaked structure have been deposited in the PDB as entries 8EBI, 8EBR, 8EC4 and 8ECF respectively.

